# Neural coding of arm and hand actions is spatially organized in motor cortex

**DOI:** 10.64898/2026.05.02.722416

**Authors:** Nicholas G. Chehade, Omar A. Gharbawie

**Author notes:** Corresponding author: Omar A. Gharbawie, Systems Neuroscience Center, University of Pittsburgh, 3501 Fifth Ave., 4069 BST-3, Pittsburgh, PA, 15261.

## Abstract

Primate motor cortex (M1) contains distinct zones with dense projections to the spinal circuits of arm and hand muscles making them critical substrates for manual behavior. But how does neural activity encoding manual actions map in space and time onto these zones? We addressed this question by recording single unit activity (n=1,573) throughout M1 in two macaque monkeys performing a reach-to-grasp task. Recordings were made with linear electrode arrays that were registered to detailed motor maps obtained with intracortical microstimulation. When units were grouped by somatotopic location, the time-resolved profiles from the arm and hand zones closely resembled time-lagged versions of the corresponding muscle activity. Thus, activity in both M1 zones differentiated between task phases and target objects. Unlike the arm and hand muscles, however, neural activity did not differ significantly between M1 arm and hand zones. In contrast, examining the spatial organization of neural selectivity for task phase revealed clear functional distinctions: reach-selectivity was stronger in the arm zone than in the hand zone and manipulate-selectivity was stronger in the hand zone than in the arm zone. Similarly, task condition decoding from hand zone activity was more accurate than from arm zone activity. Our findings collectively show that the encoding of reach-to-grasp movements is spatially clustered within the M1 forelimb representation and that the clusters are selective for function. These distinctions are graded rather than categorical, but they reveal tighter coupling between the spatio-temporal organization of M1 single unit activity and underlying cortical structure than generally assumed.

## INTRODUCTION

Seamless sequencing and integration of arm and hand movements enables them to operate as a functional unit. The arm transports the hand to desired endpoints, and the wrist orients the hand so that the fingers interact with objects. Motor cortex (M1) is central to controlling these actions because it is a major source of the descending projections that drive the spinal motoneurons of arm and hand muscles (Andersen et al., 1975; Dum and Strick, 1991; Strick et al., 2021). There is general agreement from intracortical microstimulation (ICMS) in macaque monkeys that the M1 motor map contains arm and hand zones that are contiguous, partially overlapping, and that each spans several millimeters (Kwan et al., 1978; Sessle and Wiesendanger, 1982; Park et al., 2001). But how does neural activity unfold in time across those anatomically defined zones to drive integrated arm and hand actions?

To build intuition for potential answers, let’s consider the spatio-temporal activity in M1 as an emergent property of the tuning and spatial organization of M1 neurons. In a system with tight coupling between structure and function, we could expect M1 single units to fire selectively to a specific movement (e.g., shoulder flexion for reaching or finger flexion for grasping; Fig 1I-J) executed as part of a larger motor behavior (e.g., feeding). We could also expect these movement-selective units to have well-defined spatial organizations. Thus, reach-selective units would be concentrated in the M1 arm zone and grasp-selective units would be concentrated more laterally in the M1 hand zone. In this framework, neural activity would peak in the M1 arm zone during reaching and then shift laterally to the M1 hand zone during grasping. In an alternative structure-function relationship in M1, however, the same unit selectivity could motivate different expectations about the spatio-temporal patterns of neural activity. For example, spatial intermixing of reach-selective and grasp-selective units would result in the neural activity peaking concurrently in M1 arm and hand zones during both reaching and grasping. Of course, a similar spatiotemporal pattern could emerge even if the constituent neurons were not movement-selective but were instead broadly tuned for both reaching and grasping.

**Figure 1.**
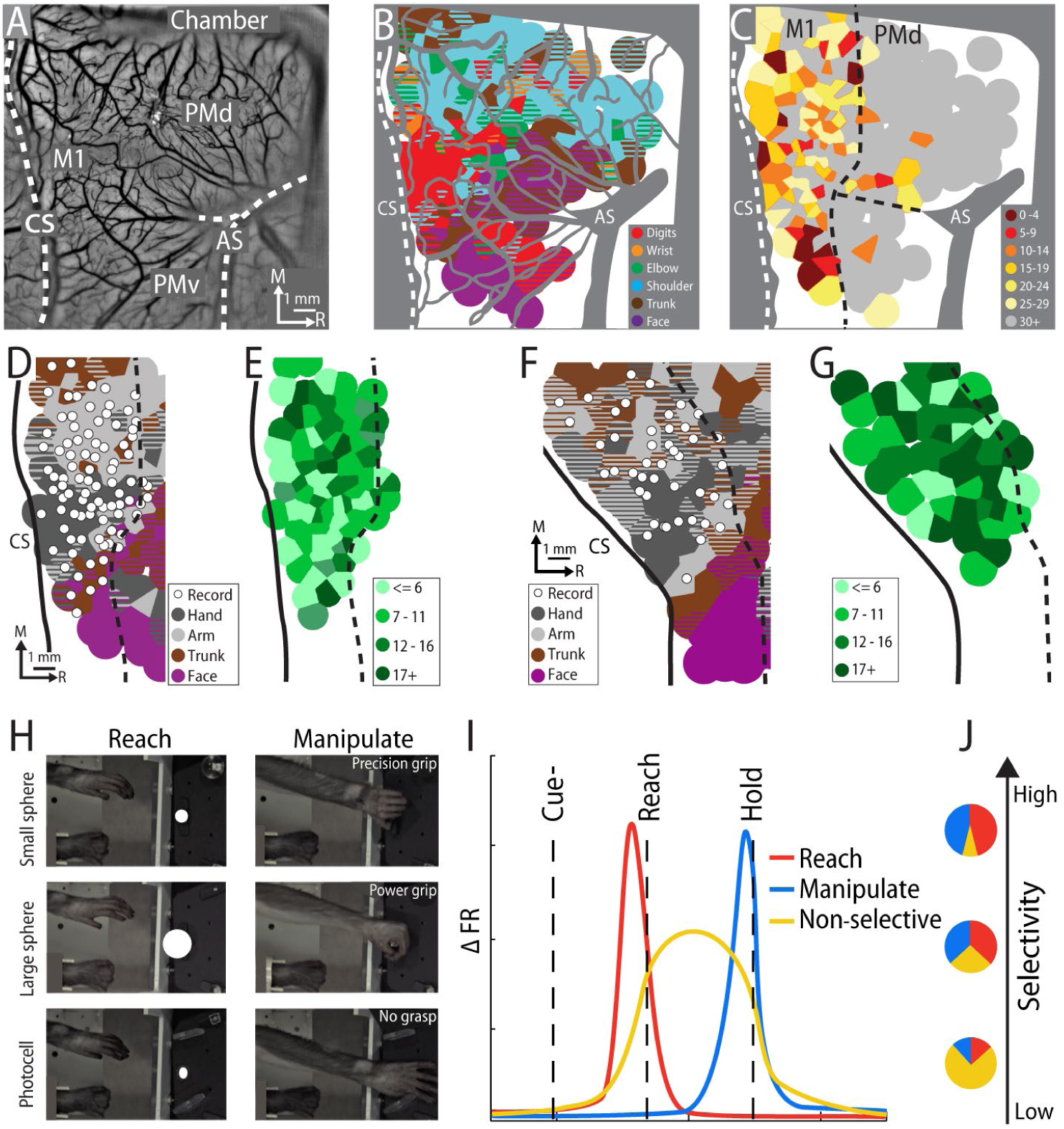
Single unit recordings across the M1 forelimb representation during reach-to-grasp task. **(A)** Image of rostral half of cranial chamber implanted in the right hemisphere (monkey G). Native dura was surgically replaced with transparent membrane. Dashed lines demarcate major landmarks: central sulcus (CS) and arcuate sulcus (AS). **(B)** Motor map based on intracortical microstimulation (217 sites). Voronoi tiles (0.75 mm radius) are color-coded according to evoked movement. Striped tiles represent dual movements. Major blood vessels and chamber edge are masked in gray. **(C)** Same as ***B*** but color-coded according to minimum current amplitude (μA). Border between M1 and premotor cortex (dashed line) drawn at the transition from low (< 30 μA) to high (>= 30 μA) current thresholds. **(D)** Same as ***B*,** but wrist and digit sites are classified as hand; shoulder and elbow sites are classified as arm. White dots are recording sites (n = 84) with linear electrode array. **(E)** Voronoi tiles of recording sites are color-coded according to number of single units (800 units). **(F**-**G)** Same as ***D-E*,** but for monkey S (53 recording sites; 795 units). **(H)** Still frames showing the reach and manipulate phases in three task conditions (rows). Task is performed with the left forelimb. Right forelimb is restrained. **(I)** Activity profile for three hypothetical types of single units. The reach-selective and manipulate-selective types are narrowly tuned for the reach and manipulate phases, respectively. The non-selective type is broadly tuned for both phases. **(J)** Pie charts reflect the proportion of each unit type at three different points along a hypothetical scale of neural selectivity for task phase.

Adjudicating between these scenarios requires precise mapping of the spatial location and tuning of every recorded unit. With these provisions, relying on previous studies to evaluate the scenarios is complicated for two reasons. First, in most M1 studies, analyses do not factor the spatial location of the units. Second, most efforts have focused on neural activity that supports arm movements (Georgopoulos et al., 1986; Schwartz et al., 1988; Kalaska et al., 1989; Moran and Schwartz, 1999; Churchland et al., 2012) or hand movements (Smith et al., 1975; Muir and Lemon, 1983; Kakei et al., 1999; Hendrix et al., 2009) but not integrated arm and hand movements. The few studies that are in that domain and that also mapped unit locations, provide different accounts on the spatio-temporal pattern of M1 activity. Some of those reports, for example, indicate that reaching and grasping movements are encoded co-extensively throughout M1 arm and hand zones (Vargas-Irwin et al., 2010; Saleh et al., 2012; Rouse and Schieber, 2016a). Neural recordings in those studies, however, sampled only a small portion (4x4 mm) of the forelimb representation in rostral M1 or sampled the entire forelimb representation but only in caudal M1. In contrast, other reports have argued in favor of a more structured spatio-temporal organization (Riehle et al., 2013; Friedman et al., 2020; Chehade and Gharbawie, 2023). For example, widefield imaging of rostral M1 showed that neural activity is spatially concentrated in patches within the forelimb representation. Follow up neurophysiology revealed that the medial patches overlapped the arm zone and were enriched with reach-selective units. In contrast, the lateral patches overlapped the hand zone and were enriched with grasp-selective units. Nevertheless, that neurophysiology dataset was relatively small (Friedman et al., 2020).

Our objective here is to evaluate the hypothetical scenarios described above, which would also shed light on the competing views from previous studies. We therefore systematically mapped the spatial organization of the time-dependent activity of M1 neurons during reach-to-grasp in two macaque monkeys. Task conditions motivated different grip types (precision-, power-, no-grip), but similar reach trajectories as objects were presented in the same location.

Neurophysiological signals were recorded with linear electrode arrays. Every penetration was made with reference to a motor map obtained from the same hemisphere with high density ICMS. This paradigm enabled us to record from hundreds of single units and to map their spatial locations and laminar depths with submillimeter resolution. From that database, we interrogated the activity profiles of the M1 arm and hand zones. Conversely, we mapped the spatial organization of neural selectivity for reach and manipulate. The results show that most task-modulated units were selective for either the reach- or manipulate-related activity and that the selectivity was organized with respect to the M1 arm and hand zones.

## METHODS

### Animals

The right hemisphere was studied in two male macaque monkeys (Macaca mulatta). Monkeys were 9-10 years old during data collection and weighed 9-11 kg. All procedures were approved by the University of Pittsburgh Animal Care and Use Committees and followed the guidelines of the National Institutes of Health guide for the care and use of laboratory animals.

### Head post and recording chamber

After an animal acclimated to the primate chair and training environment, a head-fixation device was secured to the occipital bone and caudal parts of the parietal bone. Task training with head-fixation started after ∼1 month (monkey G) or ∼9 months (monkey S) and lasted for ∼22 months. At the end of this training period, the monkey was considered ready for neural recordings. A craniotomy was performed for implanting a chronic recording chamber (30.5 x 25.5 mm internal dimensions) over motor and somatosensory cortical areas. The chamber was secured to the skull with ceramic screws and dental cement. Within the recording chamber, native dura was resected and replaced with a transparent silicone membrane (500 µm thickness) that we fabricated from a mold. Protocols for the artificial dura have been previously described in detail (Arieli et al., 2002; Ruiz et al., 2013). The walls of the artificial dura lined the walls of the recording chamber. The floor of the artificial dura (i.e., the optical window) was flush with the surface of cortex and facilitated visualization of cortical blood vessels and landmarks (Fig 1A). The walls and floor of the artificial dura delayed regrowing tissues from encroaching underneath the optical window. Single electrodes and linear electrode arrays were readily driven through the optical window without permanent deformation.

### Reach-to-grasp task

Monkeys performed a reach-to-grasp task that was similar to others (Gardner et al., 2007; Umilta et al., 2007; Nelissen and Vanduffel, 2011; Schaffelhofer and Scherberger, 2016) and matched our previous work (Friedman et al., 2020; Chehade and Gharbawie, 2023). Monkeys were head fixed in a primate chair. The left forelimb was used, and the right forelimb was secured to the waist plate. The task apparatus was positioned in front of the animal. A stepper motor rotated the carousel between trials to present a target ∼200 mm from the start position of the left hand. The relative location of the target made it visible to the monkey and facilitated consistent reach trajectories across conditions and trials. Task instruction was provided with LEDs mounted above the target. Photocells were embedded in multiple locations within the apparatus to monitor hand position and target manipulation. An Arduino board (Arduino Mega 2560, www.arduino.cc) running a custom script (1 kHz) controlled task parameters, timing, and logged the monkey’s performance on each trial. The task involved four conditions presented in an event-related design (1 successful trial/condition/block). Condition order was randomized across blocks.

***1. Two reach-to-grasp conditions.*** In a successful trial, the animal had to reach, grasp, lift, and hold a sphere. The small sphere condition (12.7 mm diameter) and the large sphere condition (31.8 mm diameter) were used to motivate precision and power grips, respectively (Fig 1H). Both spheres were attached to rods that moved in a vertical axis only. Task rules were identical for both conditions and are therefore described once. To initiate a trial, an animal placed its left hand over a photocell embedded in the waist plate. Covering the photocell for 300 ms turned on an LED, which signaled the start of the trial. Holding this start position for 5000 ms triggered the cue, which was a blinking LED. The animal had 2400 ms (monkey G) or 2550 ms (monkey S) to reach, grasp, and lift the sphere; time limits were also set for each phase. Lifting the sphere by 15 mm turned the blinking LED solid, which signaled the beginning of the hold phase. Maintaining the lifted position for 1000 ms turned off the LED, which instructed the animal to release the object and withdraw its hand back to the start position within 900 ms. Maintaining the start position for 5000 ms triggered a tone and LED blinking. After an additional 2000 ms in the start position the trial was considered successful; tone and LEDs turned off, and water reward was delivered. The animal could not initiate a new trial for another 3000 ms. Failure to complete any step within the allotted time window resulted in an incorrect trial signaled by a 1500 ms tone and a 5000 ms timeout in which the apparatus was unresponsive to the monkey’s actions. After the timeout, a new trial could be initiated with hand placement in the start position. Across both monkeys, the median failure rate per session was 15% (IQR = 6-32%) in the precision grip condition and 10% (IQR = 5-23%) in the power grip condition.
***2. Reach-only condition.*** The target was a photocell embedded into the surface of the carousel (Fig 1H). The photocell was visible to the monkey but was not graspable. The Go Cue was the same as the one used in the reach-to-grasp conditions, but here it prompted the monkey to reach and place its hand over the photocell. The hand had to cover the photocell for ≥220 ms (monkey G) or ≥320 ms (monkey S). All other task rules and steps were identical to the reach-to-grasp condition. Across both monkeys, the median failure rate per session was 19% (IQR = 7-40%).
***3. Withhold condition.*** In a successful trial, the monkey had to maintain its hand in the start position for ∼10 s. Trial initiation was identical to the other conditions. Holding the start position for 5000 ms triggered the Withhold Cue, which was distinctly different from the Go Cue in the movement conditions. Maintaining the start position for another 2800 ms triggered a tone and LED blinking. After an additional 2000 ms in the start position the trial was considered successful and rewarded. Removing the hand from the start position at any time resulted in an incorrect trial and the same consequences described in a failed reach-to-grasp trial. Across both monkeys, the median failure rate was 2% (IQR = 0-5%).

To relate the recorded neural activity to behavior, we focused on specific phases of the movement conditions. (1) *Baseline*: -4000 to -1000 ms from cue onset. (2) *Cue:* 0 to +200 ms from cue onset. (3) *Reach:* -150 to +50 ms from reach onset. (4) *Manipulate:* -250 to -50 ms from target hold onset. (5) *Withdraw:* -150 to +50 ms from withdraw onset. We limited the phases to 200 ms windows to minimize overlap between the reach and manipulate phases.

### Muscle activity

Electromyography (EMG) was recorded from 7 forelimb muscles as readout of arm and hand activity. EMG was recorded concurrently with neural activity or in separate sessions. After head fixation, the monkey was lightly sedated with a single dose of ketamine (2-3 mg/kg, IM). Sedation was confirmed from a reduction in voluntary movements with the working forelimb. At that point, pairs of stainless-steel wires (27 gauge, AM Systems) were inserted percutaneously into each muscle (∼15 mm below skin). We targeted three arm muscles: deltoideus, triceps brachii, and biceps brachii, and four extrinsic hand muscles: extensor carpi radialis (ECR), flexor carpi radialis (FCR), extensor digitorum (EDC), and flexor digitorum superficialis (FDS). The task started 45-60 min after sedation. The monkey was fully alert by that point and showed no lingering effects of sedation. The non-working forelimb was restrained and therefore could not tamper with the EMG wires.

EMG signals were filtered (bandpass 15-350 Hz) and recorded at 2 kHz with the same Ripple Neuro system that we used for neural signal acquisition. Recorded signals were segmented into trials, and their power spectral density was estimated with a discrete Fourier transform (MATLAB *fft* function, Natick, MA). Trials with power >7 μV^2^ in the 1-14 Hz range were presumed to have artifact and were excluded from further analysis. EMG signals were rectified and smoothed with a 150 ms sliding window (MATLAB *filtfilt* function). The processed traces were then aligned to 3 behavioral event times: reach onset, hold onset, and withdraw onset. To facilitate comparisons between muscles and averaging across sessions, EMG trials were normalized within a session. Thus, trials from a muscle were normalized to the peak activity of that muscle, which was the mean of the three largest peaks in that session. EMG trials from all sessions were then collated into two categories: arm or hand (2,482-2,938 trials/arm muscle; 2,482-2,556 trials/hand muscle). Within a category, we averaged one trial from each muscle to create aggregated *arm trials* and *hand trials* (Fig S1).

### Motor mapping

We used intracortical microstimulation (ICMS) to map the somatotopic organization of frontal motor areas. In monkey S, all sites (n=158) were investigated with a microelectrode in dedicated motor mapping sessions. We used the same approach in >50% of the sites (n=118) in monkey G. The remaining sites (n=99) were mapped with a linear electrode array at the end of recording sessions. In the dedicated motor mapping sessions, the monkey was head-fixed in the primate chair and sedated (ketamine, 2-3 mg/kg, IM, every 60-90 minutes). This mild sedation reduced voluntary movements but did not suppress reflexes or muscle tone.

A hydraulic microdrive (Narishige MO-10) connected to a customized 3-axis micromanipulator was attached to the recording chamber for positioning a tungsten microelectrode [250 μm shaft diameter, impedance = 850 ± 97 kΩ (mean ± SD)] or a platinum/iridium microelectrode [250 μm shaft diameter, impedance = 660 ± 153 kΩ (mean + SD)]. A surgical microscope aided with microelectrode placement in relation to cortical microvessels. The microelectrode was in recording mode at the start of every penetration. The voltage differential was amplified (10,000×) and filtered (bandpass 300–5000 Hz) using an AC amplifier (Model 2800, AM Systems, Sequim, WA). The signal was passed through a 50/60 Hz noise eliminator (HumBug, Quest Scientific Instruments Inc.) and monitored with an oscilloscope and a loudspeaker. As the electrode was lowered, the first evidence of neural activity was considered 500 μm below the pial surface.

The microelectrode was then switched to stimulation mode and the effects of ICMS were evaluated at ≥4 depths (500, 1000, 1500, 2000 μm). Microstimulation trains (18 monophasic, cathodal pulses, 0.2 ms pulse width, 300 Hz) were delivered from an 8-Channel Stimulator (model 3800, AM Systems). Current amplitude, controlled with a stimulus isolation unit (model BSI-2A, BAK Electronics), was increased until a movement was evoked (max 300 μA). The stimulation threshold for each depth was the current amplitude that evoked movement on 50% of stimulation trains.

One experimenter controlled the location and depth of the microelectrode. A second experimenter, blind to microelectrode location, controlled the microstimulation. Both experimenters inspected the evoked response and discussed their observations to agree on the active joints (i.e., digits, elbow, etc.) and movement type (flexion, extension, etc.). Movement classification was not cross-checked against EMG recording or motion tracking. The overall classification for a given penetration included all movements evoked within 30% of the lowest threshold across depths. The location of each penetration (500-1000 μm apart) was recorded in relation to cortical microvessels. Color-coded maps were generated from this data using a voronoi diagram (MATLAB *voronoi* function) with a maximum tile radius of 900 μm (Fig 1B). The rostral border of M1 was marked to separate sites with thresholds <30 μA from higher threshold sites (Fig 1C). Motor maps were further simplified by consolidating site classifications into broader categories (e.g., elbow and shoulder became arm; Fig 1D,F).

The mapping protocol was similar for penetration sites stimulated with a linear electrode array (32 or 24 channels, 15 μm contact diameter, 100 μm inter-contact distance, 210-260 μm probe diameter; V-Probe, Plexon). Each penetration was mapped ∼2.5 hours after the linear array was inserted into cortex, which was also the end of the neural recordings during task performance. Only 1 penetration was mapped per session. Microstimulation parameters were identical to the ones used with the microelectrode but were controlled here using a Ripple Neuro (Scout model, Salt Lake City, UT). Channels were stimulated one at a time and every other channel was used. In this setup, one experimenter controlled the microstimulation and classified the evoked movements.

### Neural recordings

We recorded neurophysiological signals from M1 zones that are on the dorsal surface of the precentral gyrus (i.e., old M1). In every recording session, a linear electrode array was acutely inserted orthogonal to the cortical surface. Two types of arrays were used. (1) V-probe: 32 or 24 channels, 15 μm channel diameter, 100 μm inter-channel distance, 210-260 μm shaft diameter (Plexon Inc). (2) LMA-v2: 32-channel, 12.5 – 15 μm channel diameter, 100 μm inter-channel distance, 300 μm shaft diameter (Microprobes). The position of the array was controlled with a custom 3-axis micromanipulator outfitted with a hydraulic microdrive (Narishige MO-10). We used a surgical microscope to guide each penetration in relation to cortical microvessels, which were landmarks for registering recording sites to the motor map (Fig 1 D,F). At the first sign of neural activity in the deepest recording channel, we relied exclusively on the hydraulic microdrive to advance the linear array. Once ∼90% of channels were in cortex, we allowed ∼45 minutes for the array to settle. The monkey sat quietly in the dark during that period and minor adjustments were made to the depth of the array to optimize single unit isolation across channels. Raw signals were filtered (bandpass 0.3 – 7,500 Hz) and recorded at 30 kHz with a Ripple Neuro system (Scout model, Salt Lake City, UT). In a separate data stream, raw signals were filtered (bandpass 250 – 7,500 Hz) for spike extraction. A threshold (typically 2-3 σ) was manually set for each channel and distinguishable waveforms were flagged as units using time-amplitude window discriminators. Spike waveforms, spike times, and event-times were recorded at 30 kHz. Throughout each recording, micro adjustments were made with the hydraulic microdrive to limit waveform drift across recording channels.

### Spike sorting

Spike waveforms were sorted in Offline Sorter (Plexon Inc). We used two criteria to define a single unit. (1) Peak-to-trough amplitude of the average waveform was >5x the mean voltage of the entire recording from the same channel. (2) Violations of the inter-spike interval (1.8 ms) were <0.5% of the waveforms assigned to a unit. Only single units (n=1,573) were considered for analyses. Those units were recorded from 136 linear array penetrations (84 from monkey G; 52 from monkey S) that we placed in M1 arm, hand, and trunk zones. For spatial reference, the penetration locations and their affiliate single units are shown in relation to the motor maps (Fig 1D,G).

### Spike density function

Only successful trials were analyzed. For a given unit, the spike times in a trial were aligned to the onset of three behavioral events: reach, hold, and withdraw. Spikes were then counted in 10 ms bins from the start to the end of each trial. Spike counts were smoothed with a gaussian kernel density (150 ms width) to estimate the spike density and instantaneous firing rate. Each trial was then normalized by its *baseline* (-4000 to -1000 ms from cue) activity. Normalized trials became the inputs for subsequent analyses and unit classifications.

### Task modulated units

A unit was considered task modulated if its firing rate in the movement phase (+200 ms from cue to hold) differed significantly from its baseline firing rate (paired t-test, p<0.05). Subsequent analyses are limited to task modulated units (n = 1,357; 86% of single units).

### Phase selective units

Every task modulated unit was further classified according to its selectivity, if any, for the reach or manipulated phases. Thus, for every trial, we calculated the mean firing rate of the reach phase and of the manipulate phase. We then directly compared the two distributions (paired t-test). Units with significantly (p<0.05) higher firing rates in the reach phase were classified as reach-selective. Units with significantly higher firing in the manipulate phase were classified as manipulate-selective. Units with no statistical difference were classified as non-selective. Task phase selectivity was assessed independently for each movement condition. We then generated average spike density functions for the reach-selective units, manipulate-selective units, and the non-selective units.

### Selectivity index

For every task-modulated unit, we calculated a selectivity index (SI). Our objective was to report on a ratio scale a unit’s preference, if any, for the reach or manipulate phases:

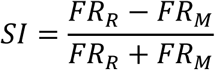

FR_R_ is the mean firing rate in the reach phase. FR_M_ is the mean firing rate in the manipulate phase. The range of SI values is +1 to -1, where +1 indicates firing exclusively in the reach phase and -1 indicates firing exclusively in the manipulate phase. A value of 0 indicates equal firing in the reach and the manipulate phases.

The SI of every unit was calculated independently for each condition. Input values were calculated from the average spike density function. The start and end times of each phase were determined from averaging those times across the trials of a given unit.

### Spatial alignment

We established a Cartesian coordinate system to co-register results across monkeys. We fit an imaginary line to the central sulcus and considered it the zero line for the rostro-caudal axis. At the border of the arm and trunk zones, we drew a line orthogonal to the one fitted to the central sulcus. The new line became the zero line for the medio-lateral axis. Thus, the intersection point of both lines was coordinate [0,0] from which we measured the spatial location of every recording site.

### Decoding

We investigated the extent to which the three movement conditions could be decoded from unit activity. We were particularly interested in relating classification accuracy to features of the recorded units like somatotopic location, laminar depth, and unit type (i.e., task phase selectivity). Unit activity in each task phase was modeled using a Poisson distribution with the λ parameter calculated from the average spike count in a task phase (Shadlen and Newsome, 1998). Thus, in each phase, the likelihood of a unit spiking 𝑥𝑥 times was calculated as:

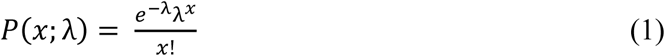

We used Naïve Bayes classifiers to predict task conditions from the spike counts of individual units (Lange et al., 2023). The principle underlying Bayesian inference here is that each instance of condition classification was determined from the maximum a posteriori probability among conditions. The posterior probability of each condition is the product of the prior probability and the marginal likelihood that the observation occurs for each condition. In the context of our model, we calculated the posterior probability ([𝑃(𝑐|𝑥)] of a given trial with 𝑥 observed spikes corresponding to a given condition (𝑐) as proportional to the product of the prior probability [𝑃(𝑐)], and the likelihood that 𝑥 spikes were observed for the condition 𝑃(𝑥|𝑐) (from Equation 1). From the set of conditions (𝐶), the posterior probability of observing 𝑥 spikes in a given condition (𝑐):

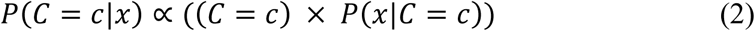

Note, the prior probability in our model was uniform across conditions (i.e., equal number of trails across conditions). Prior probability would therefore not add meaning here. To be comprehensive, the prior probability was included in Equation 2 but was then dropped in subsequent equations. After computing the posterior probabilities for each condition, we assigned the classification to the movement condition with the highest probability.

***1. Classification with one unit*.** To provide intuition for classifier training and testing, we first describe the process for one unit. For each task condition, we generated 10 sets of trials (i.e., 10-fold cross-validation). Each fold consisted of 20 trials/condition that were randomly selected with replacement from all available trials. The 20 trials were then randomly assigned to the training set (n=18) or the testing set (n=2). The training trials were averaged to obtain an expected spike count (λ) for modeling the Poisson distribution for each condition. The 10-fold cross-validation was adopted to control variance from (1) trial selection for a fold, (2) trial assignment to training and testing sets. Next, the expected spike count distributions of each condition were used to calculate the posterior probability of the 6 test trials (3 conditions x 2 trials/condition). The condition that returned the maximum posterior probability was considered the classification of the test trial. This procedure was repeated for the 6 test trials to determine the classification accuracy of a fold. The mean accuracy of 10 folds was considered the classification accuracy for a single iteration. Five hundred iterations were performed to obtain a bootstrapped distribution of classification accuracy to control variance from random selection of units as inputs to the classifier.
***2. Classification with multiple units.*** We scaled up the classification procedure so that it considered inputs from unit-sets of various sizes [1, 2, 5, 10, 15, 25, 35, 50, 75, 100]. Unit responses were modeled as probability distributions from the exponential family (e.g., Poisson) so that conditional independence between units could be assumed (Ma et al., 2006). Conditional independence gave the Bayesian classifier its Naïve quality because input from one unit was independent from all other units. As such, each unit had its own set of posterior probabilities for each task condition. Because the classifier assumed conditional independence between units, the posterior probabilities were simply the product of posterior probabilities across units. Thus, we simplified Equation 2 by removing the uniform prior (constant) and calculated the posterior probability for a given condition that considers 𝑁 units as:

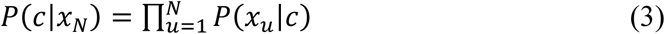

Note that unit responses were pooled from different recording sessions and monkeys. For a given test trial, we computed independent posterior probabilities for each unit. The posterior probability for a condition was estimated from the product of each unit’s (𝑁) posterior probabilities for that condition. Thus, each unit ‘voted’ and the vote was weighted according to the likelihood that the current observation corresponded to the respective condition. The predicted condition was the one that yielded maximum posterior probability.

### Statistical Analyses

We used paired t-tests (MATLAB) on every unit determine if it was task modulated and to determine if it was selective for task phase. In those tests, trials supplied the distributions of values analyzed. We used repeated measures ANOVA (SPSS) to determine whether unit activity differed across somatotopic zones, laminar depth, unit type, task phases, and task conditions.

Somatotopic zones, laminar depths and unit types were treated as between-subject factors, whereas task phases and conditions were treated as within-subject factors. For these analyses, the average spike density functions of the individual units supplied the distributions of values analyzed. Post-hoc tests were corrected for multiple comparisons (Bonferroni).

## RESULTS

Two monkeys performed an instructed forelimb task that consisted of four conditions: (1) reach-to-grasp with precision grip, (2) reach-to-grasp with power grip, (3) reach-only, and (4) withhold (Fig 1H). Both monkeys completed a minimum of 20 successful trials/condition/session (median [IQR] = 60 [59-60] trials for monkey G; 31 [30-33] trials for monkey S). In the hemisphere contralateral to the working forelimb, we used ICMS for high density mapping of the M1 motor outputs. Motor map organization was consistent across monkeys. The forelimb representation occupied most of the mapped territory and was flanked medially by trunk zones and laterally by neck and face zones (Fig 1D,F). Within the M1 forelimb representation, the main hand zone (i.e., digits and wrist) was surrounded by an arm zone, or an arm and trunk zone. This nested organization is consistent with previous M1 maps obtained in macaques with ICMS trains (Murphy et al., 1978; Sessle and Wiesendanger, 1982) and stimulus triggered averaging of EMG (Park et al., 2001), and also consistent with human maps from fMRI (Meier et al., 2008).

We leveraged the microvessel patterns visible through the artificial dura (Fig 1A) to target sites within the motor map for neurophysiological recordings (Fig 1D,F). To that end, we placed linear electrode arrays in 136 acute penetrations (monkey G= 84, monkey S=52; 1 penetration/session; 32-channel arrays in most penetrations). We recorded 1,573 well-isolated single units (monkey G= 895, monkey S= 678). The number of single units varied across penetrations (median = 10 [5-15] units; Fig 1E,G). Waveforms from two representative single units are shown in Fig 2A. Most units (n = 1,357; 86% of single units) modulated their activity during movement in at least one task condition and were therefore considered task modulated.

**Figure 2.**
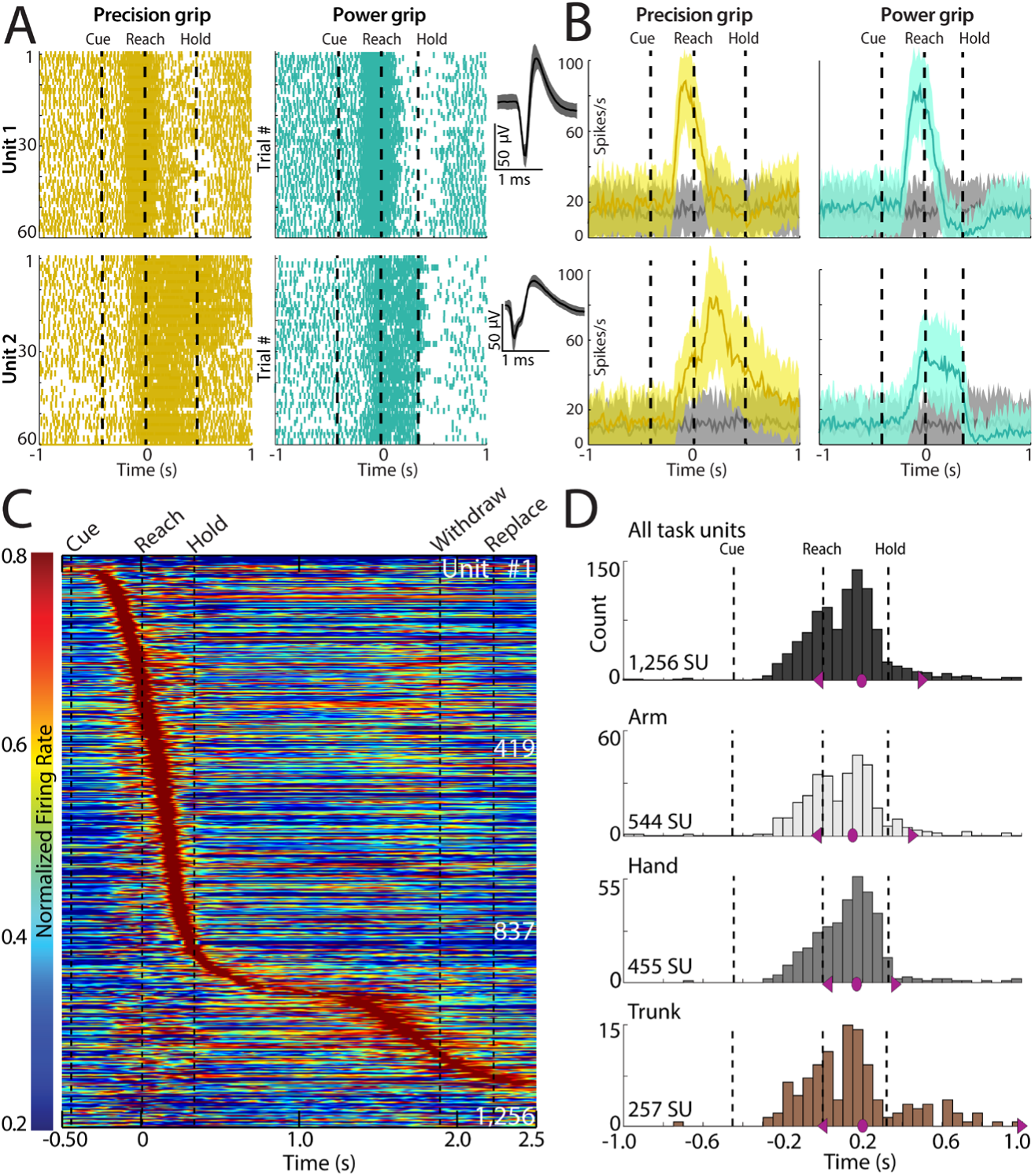
Peak firing for M1 units is concentrated in the manipulate phase. **(A)** Spike rasters from two units recorded on the same linear array. Rows are individual trials from the precision and power grip conditions. Time is in 10 ms bins and centered on reach onset. Insets show average waveforms (mean ± 3SD). **(B)** Peri-event time histograms (PETHs; mean ± SD) for the rasters in ***A***. Instantaneous firing rates for each trial were calculated in 10 ms and then smoothed with a gaussian kernel (bandwidth = 150 ms). Line plot colors match ***A***. Gray line plot is from the withhold condition and shown here for context. **(C)** Heat map of M1 single unit activity in the power condition (1,256 units; 2 monkeys). Each row is the average PETH from one unit after normalization by peak firing rate during movement. Units are sorted in ascending order of time of peak firing. **(D)** Counts of firing rate peaks relative to reach onset (1 count/single unit; 50 ms bins). Top histogram includes all task modulated units. The data in that histogram is subdivided by somatotopic zone in subsequent panels. Purple symbols demarcate the median and 25th and 75th percentiles.

### M1 units encode all task phases

After inspecting the raster plots and peri-event time histograms (PETHs) of all task-modulated units, we made some general observations that we describe here with two representative units (Fig 2A-B). (1) Firing rates were modulated in the movement conditions (Fig 2A-B, color) but remained at baseline in the withhold condition (Fig 2B, gray). (2) Firing rates and temporal profiles differed between units but were consistent across trials from individual units. (3) For some units, temporal profiles were consistent across movement conditions (e.g., unit 1). In other units, however, temporal profiles differed between movement conditions (e.g., unit 2). (4) In most units, firing modulation was limited to only a few hundred milliseconds of the entire movement trial. In unit 1 for example, activity increased sharply ∼200 ms after the Go Cue, peaked with reach onset, and returned to baseline before the object was grasped. The temporal profile and peak magnitude were both almost identical in the precision and power conditions. Unit 1 was therefore likely selective for reaching. Unit 2 also achieved similar peaks during the reach phase of both conditions. Nevertheless, the highest peak occurred during manipulation in the precision condition. Unit 2 was therefore likely selective for precision grasp.

For a high-level survey of task-modulated units, we examined the relationship between the timing of peak firing and the motor map. First, the average PETH of every unit was normalized by its peak. Those PETHs were then sorted by the latency of their peaks (Fig 2C). The heatmap shows that firing peaks spanned all task phases: cue onset to withdrawal onset. Most peaks, however, occurred during reach and manipulate (i.e., ∼200 ms before reach to hold).

Formal count of the number of units as a function of peak time (50 ms bins) revealed an asymmetric bimodal distribution (Fig 2D *top*). The smaller peak was within 100 ms of reach onset. The larger peak, however, was at the midpoint between reach onset and hold, which coincided with object manipulation. Parsing the unit counts by motor map location showed that the shape of the bimodal distribution was conserved in the arm and trunk zones (Fig 2D). In contrast, the distribution in the hand zone was closer to unimodal with leftward skew.

Nevertheless, the latency of peak firing did not differ statistically across somatotopic zones [ANOVA: F(2,1253) = 1.62, p = 0.20]. Thus, all zones contained units that peaked during reaching and even more units that peaked during manipulation.

### Neural activity across task conditions and phases

Next, we directly compared the temporal profiles of units from the M1 arm and hand zones. Thus, for each unit, we generated trial PETHs (10 ms bins) aligned to reach onset and then averaged across trials from the same task condition. Unit PETHs were individually normalized by their respective baselines, smoothed (150 ms gaussian kernel), and then averaged across recording locations in the arm zone or across locations in the hand zone (Fig 3A-B).

**Figure 3.**
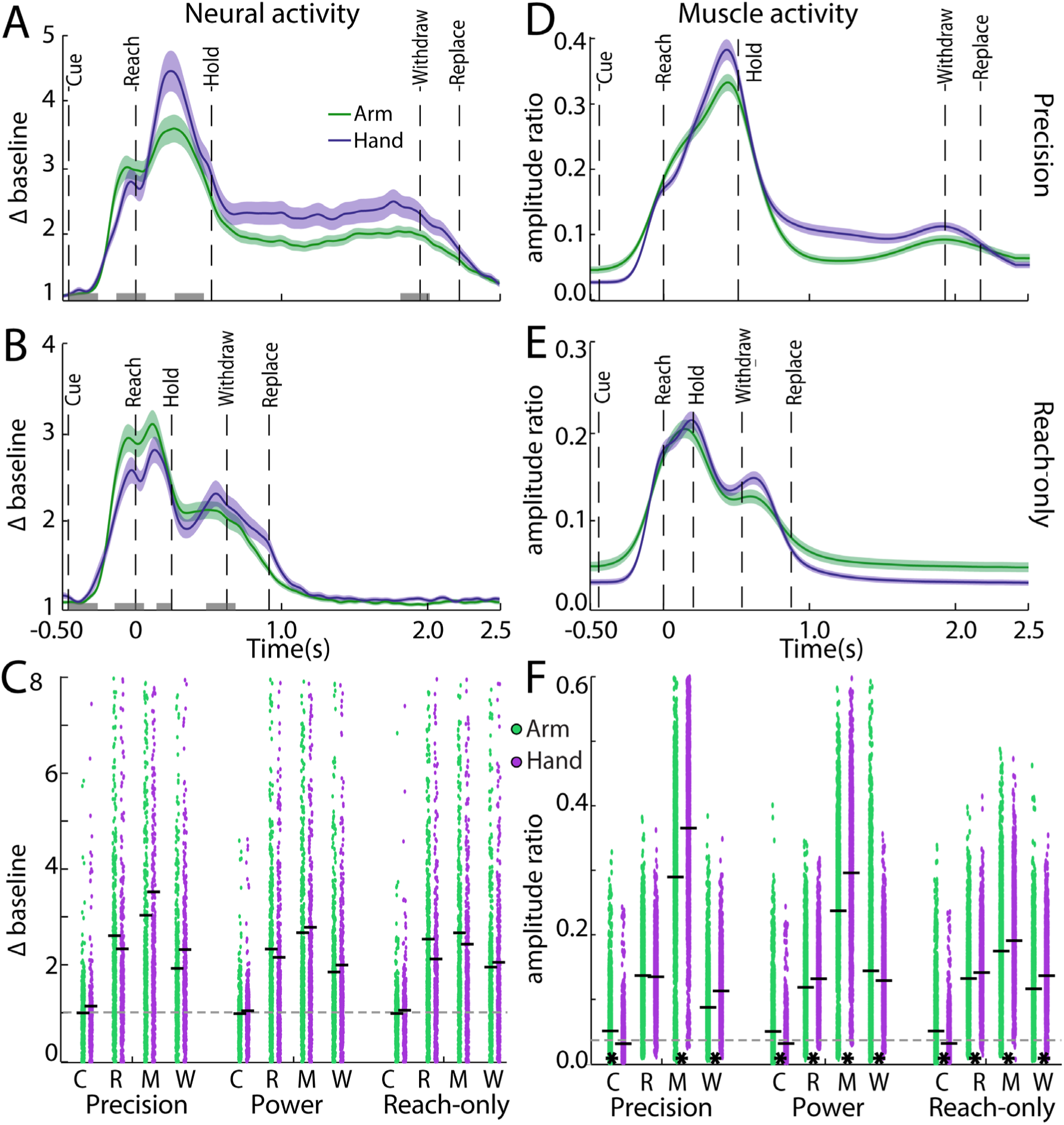
Activity is statistically distinguishable between arm and hand muscles but not between M1 arm and hand zones. **(A)** Average PETHs for single units recorded in M1 arm (588 units) and M1 hand (463 units) during the precision condition. Firing rates are expressed as multiples of baseline (mean ± SEM). PETHs are aligned to reach onset. Grey rectangles depict 4 task phases for statistical analyses (200 ms per phase). **(B)** Same as ***A***, but for the reach-only condition. **(C)** Each datapoint is from one single unit and reports its average activity in a specific task condition and task phase (C, R, M, W = cue, reach, manipulate, withdraw). Black horizontal line marks the mean of each distribution. Broken line is at baseline. **(D)** Arm and hand muscle activity in the precision condition. The arm trace is an average of pseudo-trials (mean ± SEM; 2,912 trials) based on percutaneous recordings from the deltoid, triceps, and biceps. The hand trace is an average of pseudo-trials (mean ± SEM; 2,530 trials) based on recordings from extensor carpi radialis, flexor carpi radialis, extensor digitorum, and flexor digitorum. **(E)** Same as ***D***, but the reach-only condition. **(F)** Same as ***C***, but for EMG activity. **\*** Significant pairwise differences (p < 0.001) between arm and hand muscles. All panels are based on 2 monkeys.

In the precision condition, activity in the arm zone increased sharply from ∼250 ms before reach onset to a local peak ∼100 ms before reach onset (Fig 3A). During the reach, activity settled into a saddle point and then climbed again to a global peak during object manipulation. Activity then declined to a plateau during the hold phase (i.e., hold to withdraw) and returned to baseline after the hand was withdrawn and replaced in the start position. Overall, this temporal profile is consistent with previous reports (Umilta et al., 2007; Rouse and Schieber, 2016a; Schaffelhofer and Scherberger, 2016). The average PETH from the hand zone was generally similar to the one from the arm zone but there were key differences. First, in the lead up to reach onset, activity was lower in the hand zone than in the arm zone. Second, the global peak was higher in the hand zone than in the arm zone. That separation lasted into the hold and withdrawal phases. The overall temporal profiles form both zones were consistent with the counts of peak firing (Fig 2D).

Temporal profiles from the reach-only condition resembled truncated versions of the precision condition (Fig 3A-B). This is not surprising because trials from the reach-only condition were ∼1 s shorter than those from the precision condition. Nevertheless, in the reach-only condition, activity levels were higher in the arm zone than in the hand zone during the movements leading up to hold onset. That same period also had two near-symmetric peaks, in the activity of the arm and hand zones. In contrast, in the precision condition, the equivalent peaks were strongly asymmetric and more separable between the arm and hand zones.

To quantify differences between the temporal profiles, we computed the unit firing rates in four task phases: cue, reach, manipulate, and withdraw (200 ms/epoch). That data was then analyzed with a mixed ANOVA (Fig 3C; between-subject factor: somatotopic zone; within-subject factors: task condition and task phase). Surprisingly, firing rates did not differ between M1 arm, hand, and trunk zones [F(2, 1345) = 0.87, p > 0.05]. Firing rates differed, however, between task conditions [F(2, 2690) = 14.59, p < 0.001)] and between task phases [F(3, 4035) = 196.50, p < 0.001]. Post hoc tests confirmed that firing rates were highest in the precision condition (precision > power; precision > reach-only) and differed between all pairs of task phases (manipulate > reach > withdraw > cue). Both observations are consistent with previous reports (Muir and Lemon, 1983; Umilta et al., 2007). Despite a significant 3-way interaction [task condition x task phase x somatotopic zone; (F(12, 8070) = 3.94, p < 0.001)], none of the post-hoc tests reported firing rate differences between the arm and hand zones (Fig 3C). The results collectively indicate that neural activity in both arm and hand zones was sensitive to task condition and task phase but did not differ between the two zones.

### Muscle activity across task conditions and phases

We expected the neural activity results from the arm and hand zones to show up in the behaviors of the arm and hand. To test this possibility, we first synthesized *arm trials* by averaging EMG trials from 3 arm muscles (deltoideus, triceps brachii, and biceps brachii; Fig S1). Separately, we synthesized *hand trials* by averaging EMG trials from 4 hand muscles (extensor carpi radialis, flexor carpi radialis, extensor digitorum, and flexor digitorum; Fig S1). Indeed, the temporal profiles of the synthesized trials (Fig 3D-E) closely resembled the average PETHs (Fig 3A-B). Not surprisingly, muscle activity lagged neural activity, which was most apparent in the timing of the global peak and the peak during withdrawal.

We analyzed the EMG with the same procedures used for the neural activity. Thus, for every synthesized muscle trial, activity was computed for 4 task phases. That dataset was then analyzed with a mixed ANOVA (Fig 3F; between-subject factor: forelimb segment; within subject factors: task condition and task phase). The tests reported significant effects of task condition and task phase that were consistent with the neural activity results. Nevertheless, hand muscles were generally more active than arm muscles [F(1,5589) = 115.39, p < 0.001)] but there was no equivalent difference between M1 arm and hand zones. We also found a significant 3-way interaction in the EMG [task condition x task phase x forelimb segment; F(6, 33534) = 296.06, p < 0.001]. Post hoc tests confirmed that hand muscles were more active during movement than arm muscles (Fig 3C). Overall, the results indicate that muscle activity mirrored the neural activity but that the statistical differences between arm and hand muscles were not reflected as differences between the M1 arm and hand zones.

### Higher neural modulation in superficial layers

Next, we considered the possibility that pooling units across cortical layers could have masked differences between the M1 arm and hand zones. We therefore separated units recorded on superficial channels (0-1600 μm) from units recorded on deep channels (1700-3200 μm). This rough division was intended to separate layers 1-3 from layers 5-6. Indeed, the laminar groups (Fig S2A-D) widened the gap that we observed between PETHs from the arm and hand zones (Fig 3A-B). We therefore reanalyzed the neural dataset with the same mixed ANOVA but added laminar compartment as another between-subject factor. The significant effects of task condition and phase on neural activity (Fig 3E-F) were conserved here (Fig S2E-F). The present analysis also showed that firing rates were higher for superficial units than for deep units [F(1,1342) = 4.88, p<0.05]. There was also a significant 4-way interaction [task condition x task phase x laminar compartment x somatotopic zone; F(12, 8052) = 2.52, p < 0.01]. Post hoc tests confirmed differences in three pairwise comparisons between the arm and hand zones (Fig S2E-F). Thus, separating M1 units by laminar compartment revealed functional differences between the arm and hand zones that were statistically undetectable when units were pooled across cortical layers. Nevertheless, the overall impact of the laminar separation was relatively small given that most pairwise comparison between M1 arm and hand zones were not significant.

### Spatial patterns of M1 activity across task phases

Next, we sought to examine the spatial organization of the neural activity. We therefore averaged across units recorded from the same linear array and then generated maps from that data. Figure 4A shows the progression of neural activity across task phases in the precision condition. During cue onset, activity was largely unchanged from baseline. At reach onset, however, activity increased in many recording sites in both arm and hand zones. Most of those sites remained active during object manipulation and additional sites, especially in the hand zone, increased their activity. During object hold (not shown), activity attenuated across most sites and that pattern was maintained during forelimb withdrawal.

**Figure 4.**
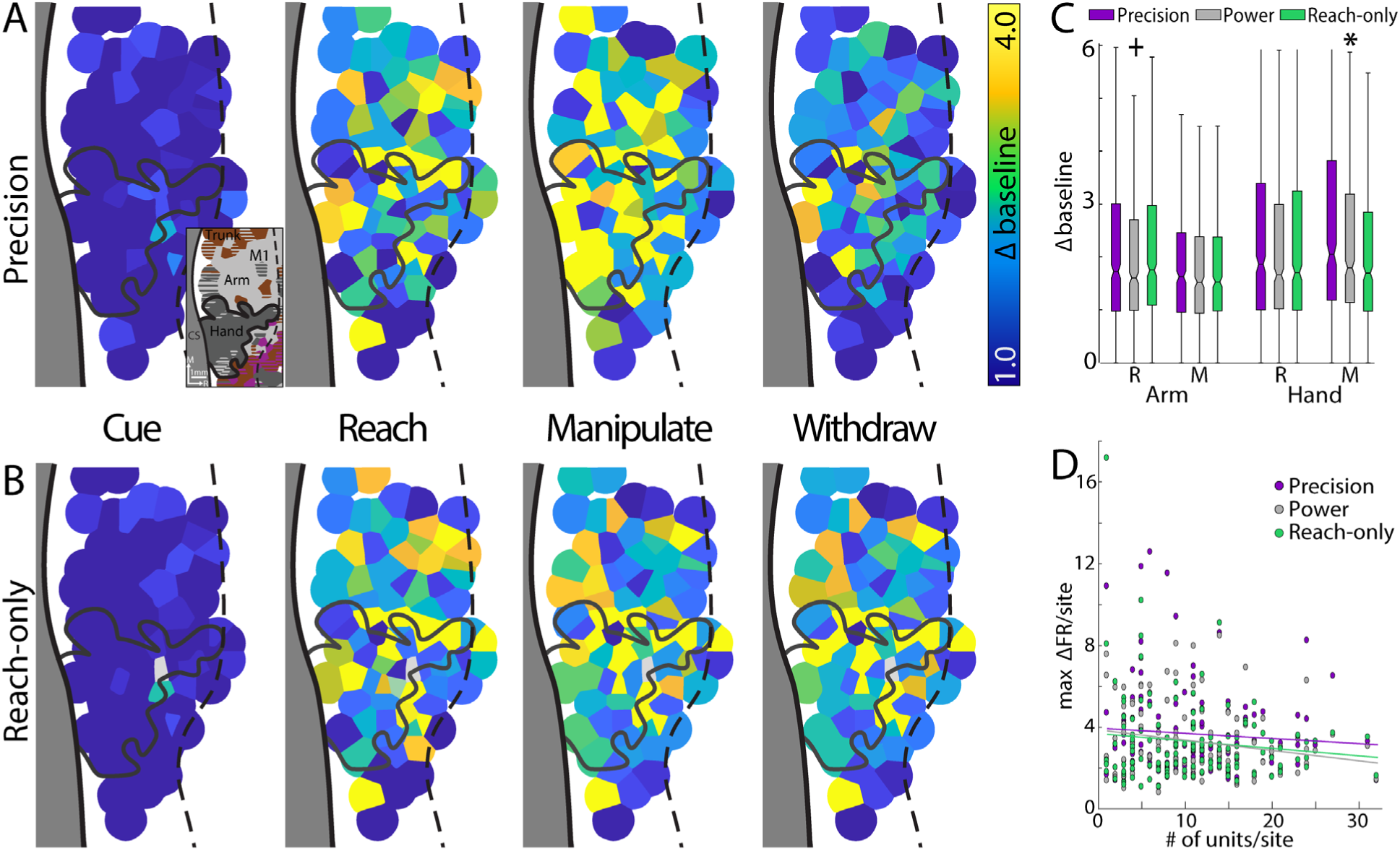
Neural activity is spatially structured in M1 across task phases. **(A)** Neural activity from the precision condition is mapped onto recording sites. Each map shows the average activity in one task phase. Activity has mosaic spatial structure and differs across the three movement phases. Voronoi tile radius = 0.75 mm. Inset shows simplified motor map for spatial reference. CS = central sulcus. M1 hand zone is outlined in black. Broken line is border between M1 and premotor cortex. **(B)** Same as ***A***, but for the reach-only condition. Here, activity patterns are qualitatively similar in the reach and manipulate phases. **(C)** Distributions of unit activity in the reach (R) and manipulate (M) phases (both monkeys). The dividing line in each box is the median. Box edges mark the 25^th^ and 75^th^ percentiles of the distribution. Whiskers span 1.5x interquartile range. Units recorded in the M1 arm zone were differentially active across conditions in the reach phase. **+** Post hoc tests indicate that power < precision and reach-only (Bonferroni corrected, p<0.05). Units recorded in the M1 hand zone were differentially active across conditions in the manipulate phase. **\*** Post hoc tests confirmed that precision > power > reach-only (Bonferroni corrected, p<0.05). **(D)** Peak firing rate from every recording site plotted as a function of number of units/site (both monkeys). Linear regressions showed no significant relationship between firing peaks and number of units (adjusted R^2^ = 0.002, -0.014, 0.021 for precision, power and reach-only, respectively).

The overall spatio-temporal pattern was similar in the reach-only condition (Fig 4B), but with some notable differences. The most apparent one was in the manipulate phase when the hand zone was more active in the precision condition than in the reach-only condition. This observation is supported by the mixed ANOVA from Figure 3. Specifically, the significant 3-way interaction [somatotopic zone x task phase x task condition; F(12, 8118) = 3.82, p < 0.001)]. Post hoc tests reported condition differences in the hand zone during the manipulate phase (precision > power > reach-only), but not during the reach phase (Fig 4C). Similarly, condition differences in the arm zone were present in the reach phase (power < precision & reach-only), but not in the manipulate phase. We interpret these results as evidence that the arm zone was more sensitive to the reach phase than the hand zone. Similarly, the hand zone was more sensitive to the manipulate phase than arm zone.

Another key observation from Figure 4A-B was that the maps from the movement phases contained many recording sites with little or no modulation. This was not simply due to sampling variations as there was no relationship between the number of units per site and firing rate modulations (Fig 4D). A more likely explanation is that the pattern reflects spatial clustering in task-related activity in M1.

### Reach and manipulate selectivity are spatially organized in M1

The spatial patterns in Figure 4 prompted us to investigate if there was spatial bias in the neural activity affiliated with the reach and manipulate phases. We therefore calculated a *selectivity index* (SI) for every task modulated unit to quantitatively describe its relative tuning for the reach and manipulate phases. SI values were averaged within recording sites and the data was used to generate maps of task phase selectivity (Fig 5A-D). To facilitate visualization, we rescaled the SI values to express selectivity for reach and manipulate as ratios of one another that sum to one.

**Figure 5.**
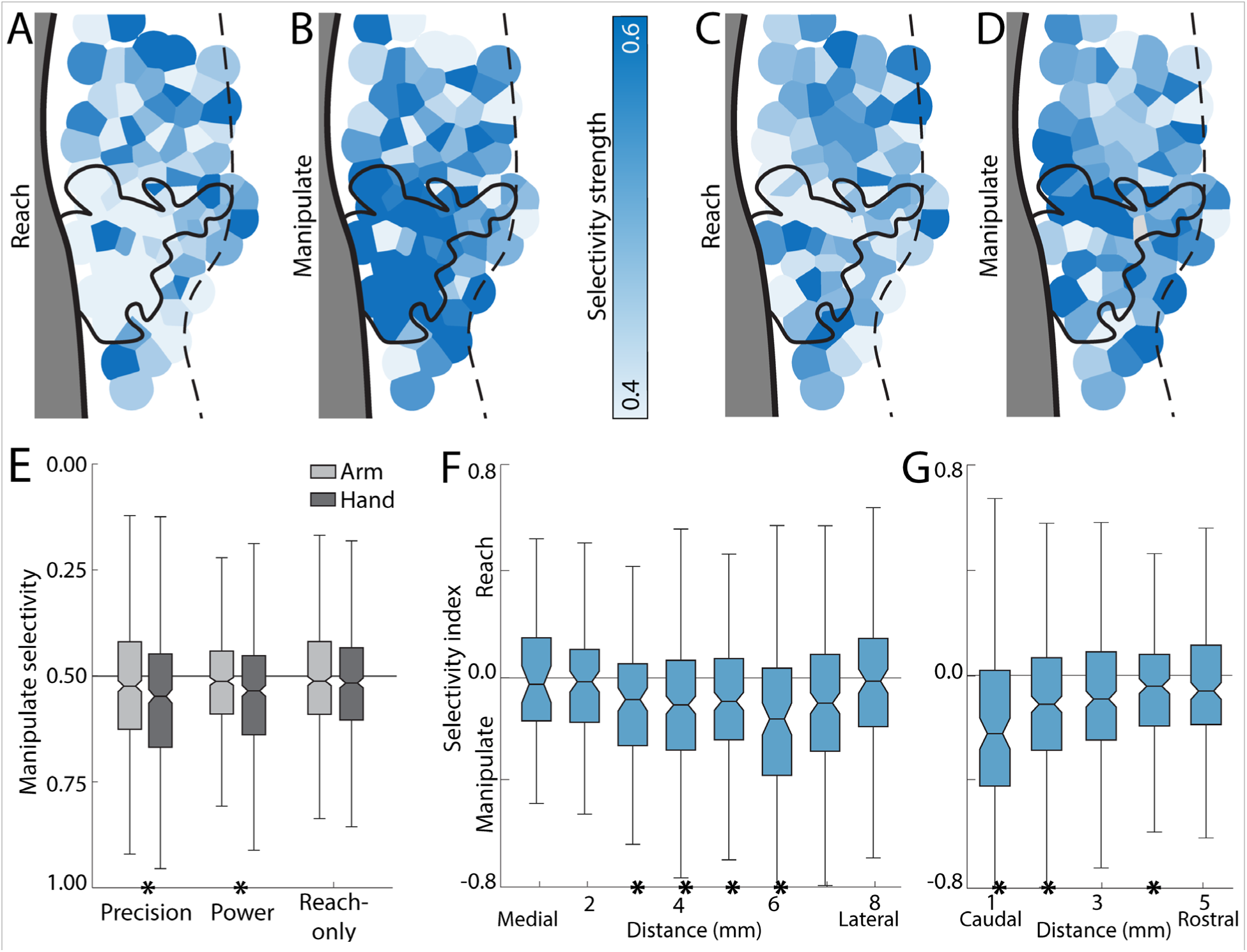
Task phase selectivity is spatially organized in M1. **(A)** Reach selectivity in the precision condition mapped onto recording sites (monkey G). Reach selectivity [reach spikes/(reach spikes + manipulate spikes)] was noticeably weak in the hand zone (black outline). **(B)** Manipulate selectivity in the precision condition. Tile colors are akin to a negative image of ***A***. Manipulate selectivity [manipulate spikes/(reach spikes + manipulate spikes)] was therefore relatively strong in the hand zone. **(C-D)** Same as ***A-B***, but for the reach-only condition. **(E)** Average manipulate-selectivity for task-modulated units in the arm and hand zones (both monkeys). **\*** p<0.01 in post hoc comparisons between arm and hand zones. **(F)** Selectivity index after binning units into 1 mm intervals in the medio-lateral axis. Data is from the precision grip condition in both monkeys. **\*** p<0.006 in one-sample t-test. **(G)** Same as ***F***, but binning here is in the rostro-caudal axis. **\*** p<0.01 in one-sample t-test. In all plots, dividing line is the median and box edges mark the 25^th^ and 75^th^ percentiles of the distribution. Whiskers span 1.5x interquartile range.

In the precision condition, the arm zone contained sites with strong reach-selectivity interspersed with sites with weak reach-selectivity (Fig 5A). In the hand zone, however, most sites had weak reach-selectivity. As can be expected from the ratio scale adopted here, the manipulate-selectivity map resembled a negative-image of the reach-selectivity map (Fig 5A-B). Thus, the arm zone contained a mixture of sites with strong and weak manipulate-selectivity, whereas most sites in the hand zone had strong manipulate-selectivity. For comparison, Figure 5C-D shows the reach- and manipulate-selectivity maps from the reach-only condition. Those maps had less contrast across sites (i.e., weaker selectivity) as compared to maps from the precision condition. To quantify the observations from Figure 5A-D, we pooled the manipulate selectivity indices across monkeys and analyzed the data with a mixed ANOVA (between-subject factor: somatotopy; within-subject factor: task condition). The analysis confirmed that manipulate-selectivity was significantly stronger in the hand zone than in the arm zone [F(2, 1337) = 5.14, p<0.01]. There was also significant interaction between somatotopic zone and task condition [Fig 5E; F(4, 2674) = 3.10, p<0.05]. Post hoc tests indicated that manipulate selectivity was stronger in the hand zone than the arm zone for the precision and power conditions but did not differ between zones in the reach-only condition.

Next, we examined the spatial organization of the SI after binning units in 1 mm increments without assignment to somatotopic zone. Our objective was to analyze the organization of SI at higher spatial resolution. To include reach- and manipulate-selectivity in the same analysis, we used the non-modified SI. Thus, +1 means neural activity was exclusively in the reach phase, -1 means neural activity was exclusively in the manipulate phase, zero means neural activity was equal in both phases. To pool data across monkeys, we co-registered their motor maps to establish a universal coordinate system [x,y] with the origin at the medial border of the arm zone and the central sulcus (Fig S3). Every recording site was then assigned to one medial-lateral interval (1 mm) and one caudal-rostral interval (1 mm). The null hypothesis is that SI values are not spatially organized. SI values within an interval should therefore average to zero. A one-sample t-test (p<0.01) compared the distribution of SI values from every interval to the null distribution (i.e., zero). In the precision condition, most intervals had distributions with negative medians (Fig 5F-G), which means that units were generally more selective for the manipulate phase than for the reach phase. The t-tests confirmed that manipulate-selectivity was significant in the middle intervals of the rostro-caudal axis and the caudal intervals of the rostro-caudal axis. Most of the significant intervals were therefore within the hand zone (Fig S3). The spatial extent of manipulate selectivity was larger in the grasp conditions than in the reach-only condition (Fig S4). In summary, Figures 5 and S4 provide evidence that selectivity for the manipulate phase was stronger in the M1 hand zone than in the arm zone.

### The temporal profiles of phase-selective units

The selectivity index results motivated us to group units by their task-phase selectivity and then reanalyze the dataset. Thus, each unit was classified as *reach-selective* or *manipulate-selective* based on a comparison (t-test) of the unit’s firing rates in the reach and manipulate phases. Units with a non-significant result were classified as *non-selective*. First, we consider the temporal profile of each unit type. In the average PETH of reach-selective units, neural activity rapidly increased from ∼250 ms before reach onset to a global peak ∼100 ms before reach onset (Fig 6A). Neural activity decreased to near baseline before hold onset and remained there until a brief bump in activity ∼100 ms before withdrawal onset. The PETH from manipulate-selective units had a distinctly different temporal profile. Here, activity climbed at a slower rate during the reach phase but then increased rapidly to a global peak in the manipulate phase (Fig 6A).

**Figure 6.**
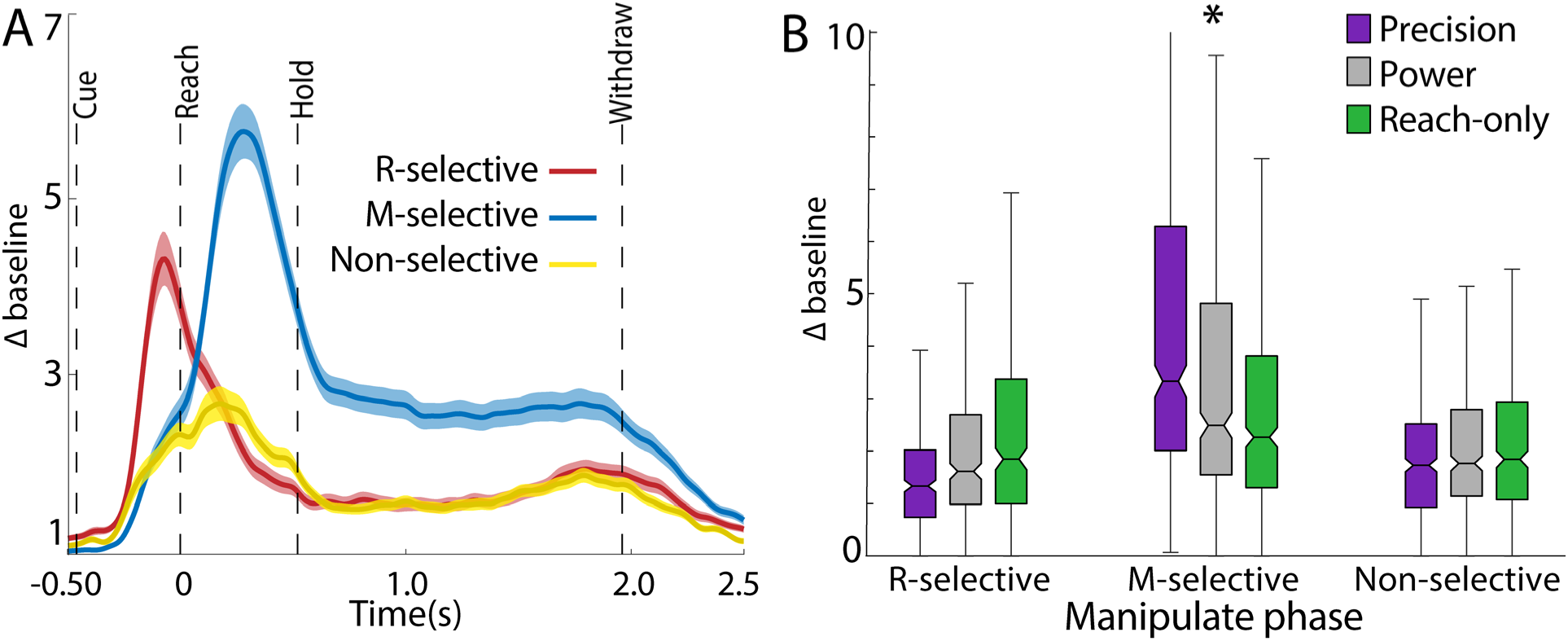
Phase selective units are tuned for task condition. **(A)** Average PETHs (mean±SEM) for the three units types in the precision condition. Units were pooled across the M1 arm and hand zones (reach-selective = 399, manipulate-selective = 498, non-selective = 402). (B) Distributions of neural activity in the manipulate phase for the three unit types. **\***Significant differences (Bonferroni corrected, p<0.05) between all pairs of task conditions in the manipulate-selective units. All panels are based on 2 monkeys.

Manipulate-selective units also sustained some level of activity throughout the hold phase. The temporal profiles of non-selective units stood out from the reach- and manipulate-selective units for having lower overall activity and a broad peak (Fig 6A).

We expanded the mixed ANOVA from Figure 3 to include unit type as a factor [between-subject factors: somatotopy, unit type; within-subject factors: task phase and task condition]. The results (main effects, interactions, and post hoc) were consistent for factors that were present in previous analyses. Thus, there was no effect of somatotopic zone on neural activity levels [F(2, 1192) = 0.22, p > 0.05], but task phase and task condition both had significant effects [F(3, 3576) = 166.11, p < 0.001; F(2, 2384) = 8.57, p < 0.001]. The new finding here was that activity levels differed between unit types [F(2, 1192) = 18.26, p < 0.001)]. Post hoc tests indicated that reach-selective and manipulate-selective units fired at higher rates than non-selective units. Moreover, there was a significant 3-way interaction [unit type x task phase x task condition; F(12, 7152) = 7.91, p < 0.001]. Post hoc tests indicated that during the reach phase, the three unit types fired differentially between the precision grip condition and at least one other task condition. In contrast, in the manipulate phase, only the manipulate-selective units fired differentially across task conditions (precision > power & reach-only; Fig 6B). Manipulate-selective units may therefore play a special role in shaping the movements needed for successful interaction with each target. Similarly, reach-selective units may play a special role in shaping movements for reaching as they were the most active unit type in that phase and the only one to differentiate between the three task conditions.

### The spatial organization of phase selective units

Next, we investigated the spatial organization of the three unit types from Figure 6. From the selectivity index results (Fig 5A-D), we expected the M1 hand zone to have a relatively high concentration of manipulate-selective units but expected a more even distribution of unit types in the arm zone. We therefore counted the number of units from each type to obtain observed distributions for each somatotopic zone (Fig 7A). For the null distribution, we assumed equal ratios across unit types (33% per unit type). We pooled the observed distributions across task conditions and compared them to the null distribution (chi square χ^2^ goodness of fit test). In the hand zone, the observed distribution differed significantly from the null distribution [χ^2^(2) = 63.09, p<0.001]. This was driven by a higher-than-expected ratio of manipulate-selective units and lower-than-expected ratio of reach-selective units. The arm zone distribution was similar to the hand zone in that it contained the same ratio of non-selective units and had a larger ratio of manipulate-selective units than reach-selective units. The ratios of the reach-selective and manipulate-selective units likely drove the significant distribution in the arm zone [χ^2^(2) = 25.40, p<0.001]. Despite the similarities between arm and hand zones, it is important to note that the arm zone had a larger ratio of reach-selective units than the hand zone. The arm zone also had a smaller ratio of manipulate-selective units than the hand zone. Finally, the trunk distribution also differed from the null distribution [χ^2^(2) = 25.40, p<0.001]. But that result was driven by a higher-than-expected ratio of non-selective units and lower-than-expected ratio of reach-selective units. Overall, the results show that M1 zones contained uneven ratios of the unit types. We interpret the high ratios of manipulate-selective units in the hand zone and non-selective units in the trunk zone as evidence for spatial organization of function in M1.

**Figure 7.**
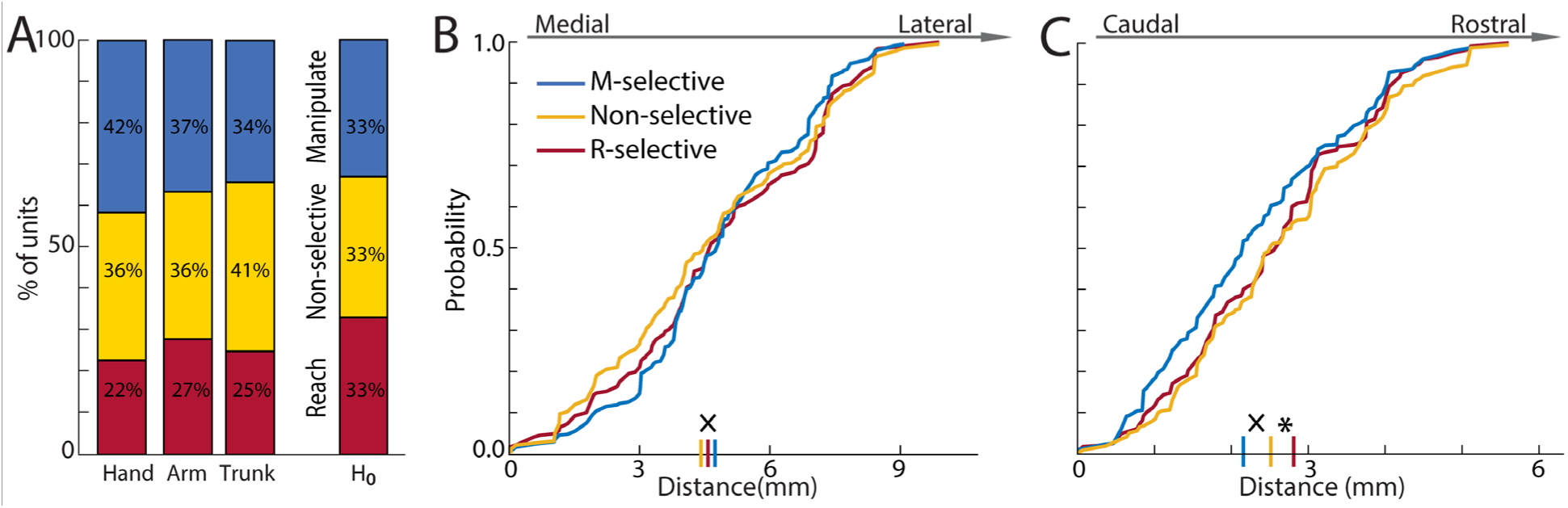
Unit types are unevenly represented in M1 arm and hand zones. **(A)** Stacked bars show the ratios of unit types in each somatotopic zones (both monkeys). Distributions reflect the mean across task conditions. *H_0_* is the null distribution with equal proportion of unit types. **(B)** The empirical cumulative distribution function (eCDF) in the medio-lateral axis of M1 for each unit type. **(C)** Same as ***B***, but in the rostro-caudal direction. **^X^** Pairwise difference (Kolmogorov-Smirnov test, p<0.05) between manipulate- and non-selective units. **\*** Pairwise difference between reach- and manipulate-selective units.

To quantify the organization of unit types at higher spatial resolution, we examined their counts as a function of their spatial coordinates (Fig S3). Thus, for each unit type, we generated an empirical cumulative distribution function (eCDF) for the M1 medio-lateral axis and a separate eCDF for the M1 rostro-caudal axis of M1 (Fig 7B-C). The eCDF reports the cumulative probability of encountering a unit type in a sweep across one of the spatial axes of M1. The probability is therefore *zero* at the start of a sweep and *one* at the end of that sweep. In general, the three unit types had overlapping probability curves in the medio-lateral axis (Fig 7B). Nevertheless, the curves were somewhat separable from 0 to 5 mm (i.e., medial half). A two-sample Kolmogrov-Smirnov test indicated that manipulate-selective units were more lateral than non-selective units [p<0.05; precision: D(452) = 0.10; power: D(442) = 0.11; reach only: D(420) = 0.10]. Separation between the probability curves was more apparent in the rostro-caudal axis (Fig 7C). Specifically, manipulate-selective units were more caudal than reach-selective units [p <0.0001; precision: D(367) = 0.20; power: D(352) = 0.18; reach-only: D(359) = 0.18] and non-selective units [p<0.001; precision: D(402) = 0.14; power: D(349) = 0.13; reach-only: D(353) = 0.14]. In summary, the cumulative distributions show that manipulate-selective units were concentrated in caudal-lateral aspects of M1, which is the location of the hand zone.

### Decoding accuracy varies across M1 populations

Our analyses thus far have been based on the same framework: grouping units according to anatomical or functional criteria then directly comparing the task-related activity of those groups. To evaluate if similar results can be obtained from a different attack angle, we measured the capacity of the unit groups to decode the three task conditions. We therefore used a Naïve Bayes classifier to decode task condition from spike counts (200 ms window). An underlying assumption here is that classification accuracy would reflect the sensitivity of the unit groups to the movement differences across conditions.

First, we investigated the effect of task phase on decoding task conditions from unit activity in M1 arm and hand zones. Classification accuracy was relatively high when task phases were matched in the training and test trials (Fig 8A, solid lines). Accuracy was near chance (33%), however, when task phases were not matched (Fig 8A, dashed and dotted lines). Training and testing on the cue phase was marginally more accurate than the results from the mismatched phases. In the top two classifiers (manipulate and reach), accuracy improved with increasing the number of units in a unit-set. We determined the numbers of units for those two classifiers to achieve 90% accuracy (Fig 8A, circles) and chose the midpoint as the unit-set size (35 units) for statistical analyses. Accordingly, we found a significant effect of task phase on classification accuracy [Fig 8B; ANOVA: F(2,1497) = 26644.40, p<0.001)]. Post hoc tests confirmed that accuracy was highest for the manipulate phase (manipulate > reach > cue; p<0.001), which is consistent with previous work (Mollazadeh et al., 2011; Schaffelhofer et al., 2015).

**Figure 8.**
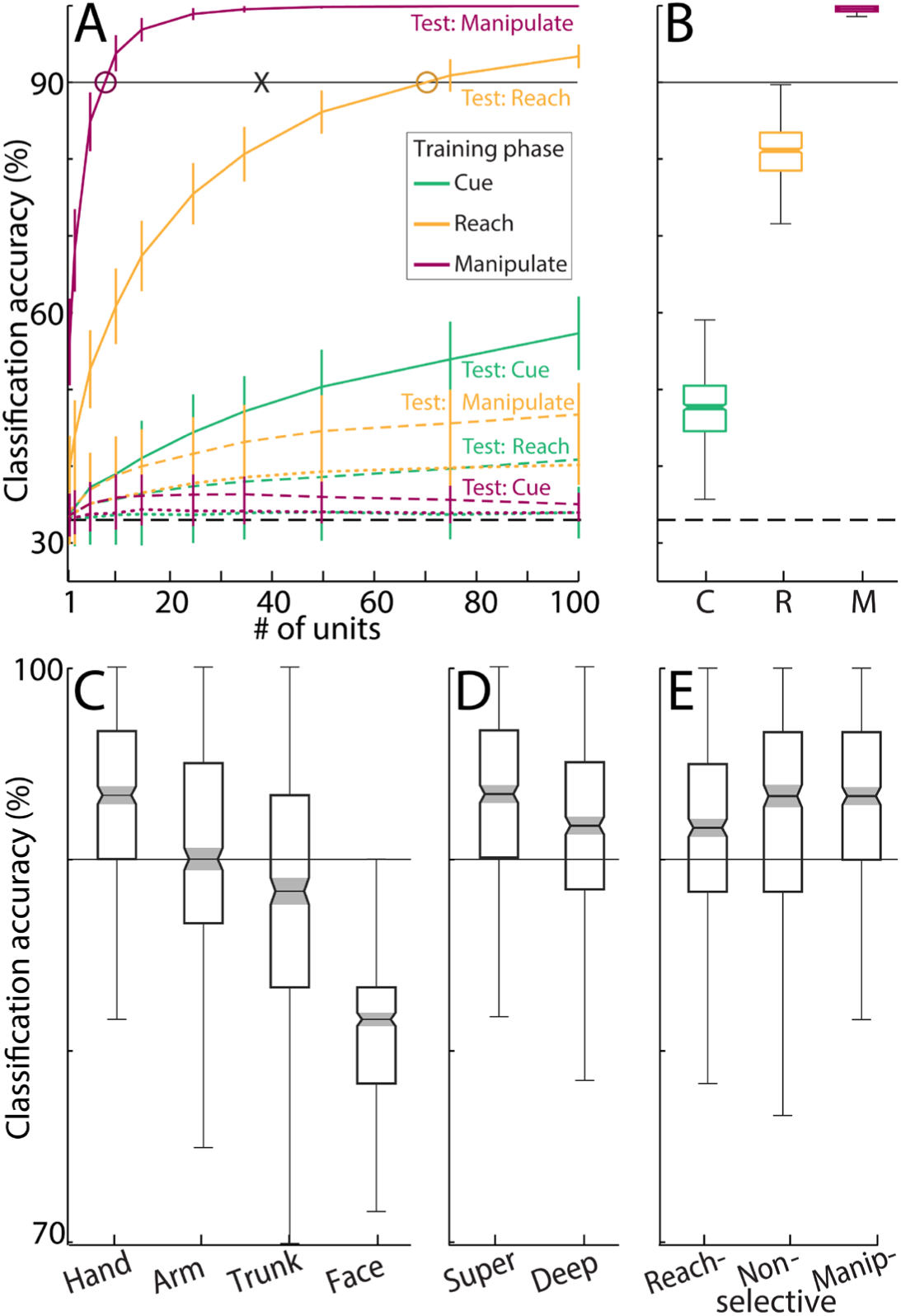
Decoding accuracy is shaped by the subpopulations of M1 units. **(A)** Accuracy of Naïve Bayes classifier plotted against number of units in the training data. Color specifies the task phase used in the training data. Solid lines indicate that task phase was matched in the training and testing data. Broken and dotted lines mean mismatch between task phase in training and testing. For each sample size, units were randomly chosen for testing the classifier and the process was repeated 500 times. Broken horizontal line marks chance-level accuracy (33%). Solid horizontal line marks 90% decoding accuracy. X: midpoint (n=38 units) for the reach and manipulate classifiers to achieve 90% accuracy (circles). Subsequent plots are based on a set sample size (35 units) randomly selected 500 times. **(B)** Classification accuracy in the three task phases (C, R, M = cue, reach, manipulate). **(C)** Classification accuracy with units grouped by somatotopic zone. **(D)** Same as ***C***, but with units grouped by laminar compartment. **(E)** Same as ***D***, but with units grouped by task phase selectivity.

The task phase results prompted us to use only the reach and the manipulate phases in subsequent analyses. We therefore averaged the accuracy from those two phases on every fold of the cross-validation. We found that somatotopic zone had a significant effect on classification accuracy [Fig 8C; ANOVA: F(3,1996) = 424.07, p<0.001]. Post hoc tests showed differences between all pairs of somatotopic zones and confirmed that classification accuracy was highest for M1 hand (hand > arm > trunk > face; p<0.001). For the final two analyses, we pooled units from the M1 arm and hand zones to test the effects of laminar compartment and unit type (i.e., selectivity for task phase). We found that classification accuracy was higher for units in superficial laminar compartments than for those in deep compartments [Fig 8D; ANOVA: F(1,998) = 7.99, p<0.005]. We also found an effect of unit type on classification accuracy [Fig 8E; ANOVA: F(2,1497) = 3.48, p<0.05]. Post hoc tests indicated that the manipulate-selective performed better than reach-selective units.

In sum, we found many differences in decoding across the M1 unit groups. Classification accuracy was highest for M1 units that were in the hand zone, or in superficial laminar compartments, or that were manipulate-selective. These observations are consistent with results obtained in our previous analyses. Thus, the anatomical and functional features that define these unit groups are potentially markers of M1 units that are sensitive to task-specific movements.

## DISCUSSION

We investigated the spatio-temporal organization of neural activity that occurs in motor cortex (M1) during manual behavior. We recorded from 1,573 well-isolated single units and registered their locations to detailed motor maps from the same hemispheres. Three findings highlight the strong relationship between neural coding of movement (function) and the M1 motor map (structure). (1) Most task-modulated units were selective for either reach or manipulation. (2) Reach- and manipulation-related activity were concentrated in patches within the M1 arm and hand zones. (3) Reach activity was stronger in M1 arm, whereas manipulation activity was stronger in M1 hand.

### The arm and hand both engage in reach and manipulation

Our EMG recordings showed that the arm and the hand were co-active across task phases. Their activity peaked during reach onset and then achieved a larger global peak during object manipulation. The magnitude of the second peak scaled with grip dexterity (precision > power > reach-only). Our EMG results were generally consistent with muscle activity and joint kinematics reported in similar tasks (Brochier et al., 2004; Rouse and Schieber, 2015, 2016b). But there was greater asymmetry here between the reach and manipulate peaks as compared to previous studies with targets presented in multiple locations. Despite the fixed target location in our task, forelimb activity differed across conditions during the reach phase and not just the manipulate phase. Similar observations have been interpreted as evidence (Paulignan et al., 1991; Haggard and Wing, 1998) against completely independent cortical networks for reaching and for grasping (Jeannerod, 1984; Gentilucci et al., 1991). Indeed, there is support from anatomical and functional studies for communication between the reach and the grasp channels of the parietal-frontal network (Cavina-Pratesi et al., 2010; Fattori et al., 2010; Davare et al., 2011; Gharbawie et al., 2011; Kaas et al., 2011).

### Evidence for spatial overlap in coding reach and manipulation

We analyzed the neural data in relation to the motor map in two fundamentally different ways. In leading with M1 structure, we grouped units by their somatotopic location and then directly compared the groups. PETHs from both arm and hand zones resembled time-lagged versions of arm and hand muscles. This fidelity suggests that muscle control was a dominant feature in the average population response from both M1 zones. Nevertheless, across movement phases, the hand was more active than the arm. That difference, however, was not reflected statistically in the neural data (Fig 3E-F).

We offer two potential explanations for the discrepancy. First, arm and hand activity was inferred from EMG recordings with acute, percutaneous electrodes. This recording modality and the anatomical separation of arm and hand muscles enabled consistently on-target measurements. Moreover, the EMG dataset was relatively large (>5,500 trials). These factors would have enhanced the likelihood of finding task-related differences between arm and hand activity. The neural dataset by contrast was “noisy”. Even though every recording site was registered with submillimeter accuracy to the motor map, specifying the somatotopic membership of M1 units was not unequivocal like defining the muscular origins of EMG traces. The uncertainty is inherent to defining motor outputs in thresholded ICMS sites. As expected, thresholding generally reduced the number of joints involved in the ICMS-evoked response despite the survival of mixed outputs in many sites (e.g., shoulder and digits). Our thresholded motor maps may have therefore underestimated the spatial overlap between arm and hand zones. A potential source of overlap is M1 columns with projections innervating the spinal circuitry of both arm muscles and hand muscles (Lemon et al., 1987; Park et al., 2004). Overlap could also reflect spatial intermingling of M1 columns with motor outputs to arm muscles and M1 columns with outputs to hand muscles (Andersen et al., 1975; Gould et al., 1986; Donoghue et al., 1992; Rathelot and Strick, 2006). In all cases, distinguishing between the neural activity of arm and hand zones becomes more challenging with increasing overlap between both zones. Second, for each M1 zone, activity was averaged across trials and aggregated across hundreds of units. This traditional approach is instructive for denoising a dataset and reducing its complexity, but it could also diminish differences that may exist between the neural populations in each M1 zone.

### Evidence for separation in coding reach and manipulation

Reversing the steps of our analysis uncovered neural activity patterns that were hidden when units were grouped by somatotopic zones. Thus, in leading with neural function, we first assessed task phase selectivity unit-by-unit and then quantified the spatial organization of the results. Units were categorized as reach-, manipulate-, or non-selective, based on direct comparison of firing rates during the reach and manipulate phases. This simple sorting system returned three distinct activity profiles (Fig 6A) that also differed from the profiles of the arm or hand zones (Fig 3A-B). A similar distinction has been reported between M1 population dynamics for reaching and grasping (Suresh et al., 2020). We conceptualize the profiles of the three unit types as the main components that shaped the average profiles of the arm and hand zones. We are formally evaluating this interpretation with dimensionality reduction (Cunningham and Yu, 2014). Our early results from principal component analysis indicate that the dominant trajectories in latent space match the temporal profiles of the three unit types.

That ∼70% of units were reach- or manipulate-selective indicates that most task-modulated units were tuned for only a subset of the movements that comprise a trial. The prevalence of task phase selectivity supports the notion that coding of reach and manipulation is separable at the single neuron level. Attributing the profiles of phase-selective units to parts of the forelimb, or a movement parameter, is complicated by the co-activation of the arm and hand across task phases. Tuning for a task phase is consistent, however, with the concept of M1 neurons encoding movement fragments (Hatsopoulos et al., 2007; Saleh et al., 2012). Fragments are movement trajectories, which are also describable in terms of temporally extensive muscle synergies (Overduin et al., 2008; Bizzi and Cheung, 2013) and can be combined to form motor actions. Although non-selective units were task modulated, they were equally active in the reach and manipulate phases. The broad, low-level activity of non-selective units suggests that they may play a more general role in the present task than phase-selective units or perhaps a role that could become more apparent with population analysis methods.

The spatial organization of the reach- and manipulate-selective units is another critical dimension in understanding the relationship between function and M1 structure. On one hand, we found that reach- and manipulate-selective units were present in both M1 arm and hand zones. This mixed organization is consistent with previous reports on spatial intermingling in M1 of reach units and grasp units (Saleh et al., 2012). It also aligns with reports on spatially co-extensive encoding of target location and intrinsic target properties in M1 (Vargas-Irwin et al., 2010; Rouse and Schieber, 2016a), dorsal premotor cortex, and ventral premotor cortex (Raos et al., 2004; Stark et al., 2007; Takahashi et al., 2017). Nevertheless, we also reported that manipulate-selective units were more prevalent in the M1 hand zone than the other two types of units. Similarly, the selectivity index tilted towards the manipulate phase in the hand zone and tilted towards the reach phase in the arm zone. These results support the notion that the neural coding of function is spatially coupled to the anatomical organization of the M1 forelimb representation. Our interpretation has support in previous findings from intrinsic signal imaging and neurophysiology (Friedman et al., 2020; Chehade and Gharbawie, 2023), BOLD fMRI (Nelissen and Vanduffel, 2011), long train microstimulation (Graziano et al., 2002; Overduin et al., 2012; Mayer et al., 2019), and 2-photon imaging (Dombeck et al., 2009).

Spatial clustering of high-activity sites (Fig 4) was another indicator of tight coupling between neural encoding of function and the M1 motor map. Our comprehensive sampling of the forelimb representation (Fig 1D-G) and the large proportion of task modulated units (86% of recorded units) convince us that spatial clustering of activity was not artifactual or incidental.

Instead, we consider clustering as evidence that the task-related movements were strongly encoded in patches within the arm and hand zones and weakly encoded in between the patches. The spatial confinement of neural modulation is consistent with findings from imaging and neurophysiology particularly in overtrained tasks (Picard et al., 2013; Friedman et al., 2020; Chehade and Gharbawie, 2023) and could reflect network efficiencies that develop with skill mastering (Peters et al., 2014). The results are also consistent with the notion of M1 containing subzones that specialize in specific motor actions (Graziano et al., 2002; Dombeck et al., 2009) and spatial heterogeneity of function in M1 (Canfield et al., 2025).

The spatial organization of function reported here was enabled by three features of our experimental design. (1) Use of high-density motor map as surrogate for M1 structure. (2) Precise registration of every recorded unit to the motor map. (3) Comprehensive sampling of single units from across the motor map (∼1500 single units from ∼130 linear arrays in 2 monkeys). We acknowledge that the structure-function relationship in sensorimotor cortex has been elegantly investigated with 2-photon calcium imaging in GCaMP-expressing mice (Salimian et al., 2025; Grier et al., 2026). The strengths of our approach must therefore be considered in the context of the investigative tools currently available in macaque monkeys and their relatively large brains.

### Insights from task condition selectivity

Sensitivity to task conditions varied between M1 zones in ways that support coupling between neural function and M1 structure. Specifically, activity levels in the M1 arm zone differed across conditions during the reach but not during manipulation. In contrast, activity in the M1 hand zone differed across conditions during manipulation but not during the reach.

Classification of task condition was also more accurate with spike counts from the M1 hand zone than with spike counts from the M1 arm zone. A similar medio-lateral gradient has been reported for decoding accuracy of imagined arm and hand movements with signals from multi-electrode arrays in human M1 (Kunigk et al., 2025). The interactions that we reported between M1 somatotopy, task phase, and task condition, collectively point to preferential coding of reach-related properties in the M1 arm zone and manipulate-related properties in the M1 hand zone.

## Conclusion

Neural activity that supports reach and manipulation is organized in M1 in both space and time. The coupling that we reported between neural coding of movement and the motor map suggests that the structure-function relationship in M1 is stronger than generally assumed. In this framework, the broader repertoire of manual actions could be spatially organized as partially overlapping zones within the M1 forelimb representation. Each zone would be enriched with neurons tuned for movements that define the actions.

## Acknowledgments

Project was supported with funds from NIH (R01 NS105697); Whitehall Foundation (2017-12-94), and University of Pittsburgh Brain Institute. We are grateful to ToniAnn Zullo and Katryna Kinnard for outstanding animal care before, after, and during, all procedures. We thank Drs Jennifer L. Collinger, Rob S. Turner, and Aaron P. Batista for many insightful discussions about the results. We thank Dr. Steven M. Chase for advice on implementation of the Naïve Bayes classifier.

## Contributions

NGC & OAG designed and performed experiments, analyzed data, and wrote the manuscript. NGC prepared all figures. OAG obtained funding.

## Data and scripts available for download

https://osf.io/wqzfg/overview

## SUPPLEMENTAL

**Figure S1.**
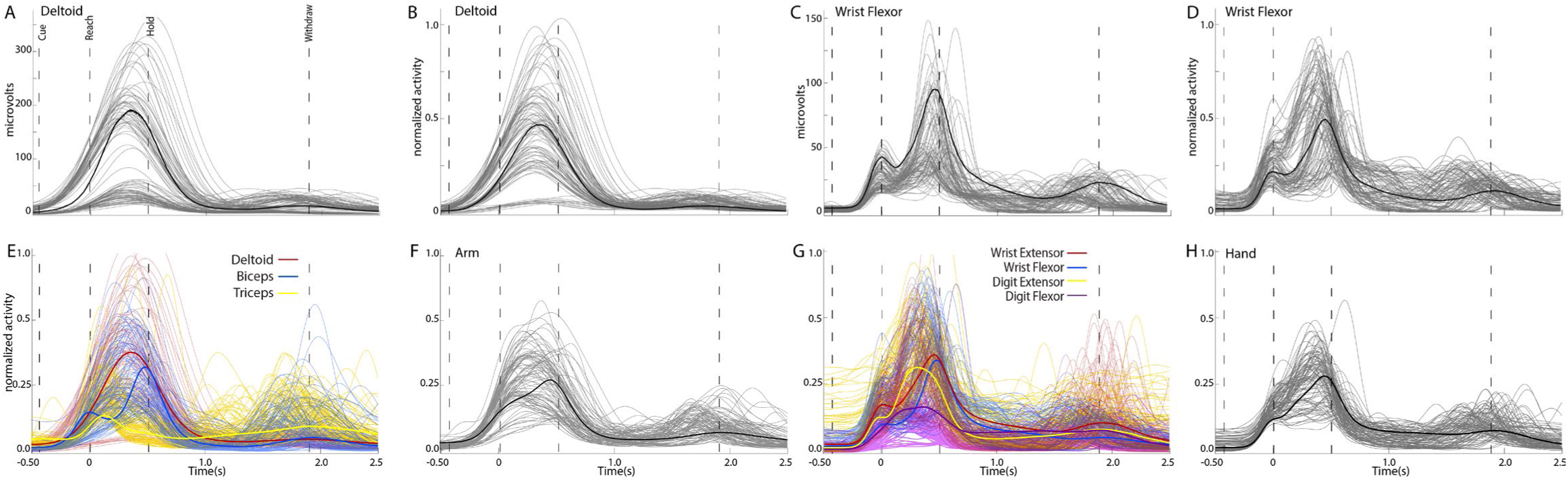
Pseudo-trials for arm and hand muscle activity. Plots are based on EMG recordings in the precision condition; 100 trials/muscle recorded over three sessions. **(A)** Deltoid trials after rectification and smoothing. Black line is the mean. **(B)** Same as ***A***, but here trials are normalized to the global peaks of their respective recording sessions. **(C-D)** Same as ***A-B***, but for the wrist flexor muscles. **(E)** Normalized trials from three arm muscles. Thicker lines are means. **(F)** Pseudo-trials generated from averaging 1 trial/muscle from ***E***. (**G-H)** Same as ***E-F***, but for four hand muscles.

**Figure S2.**
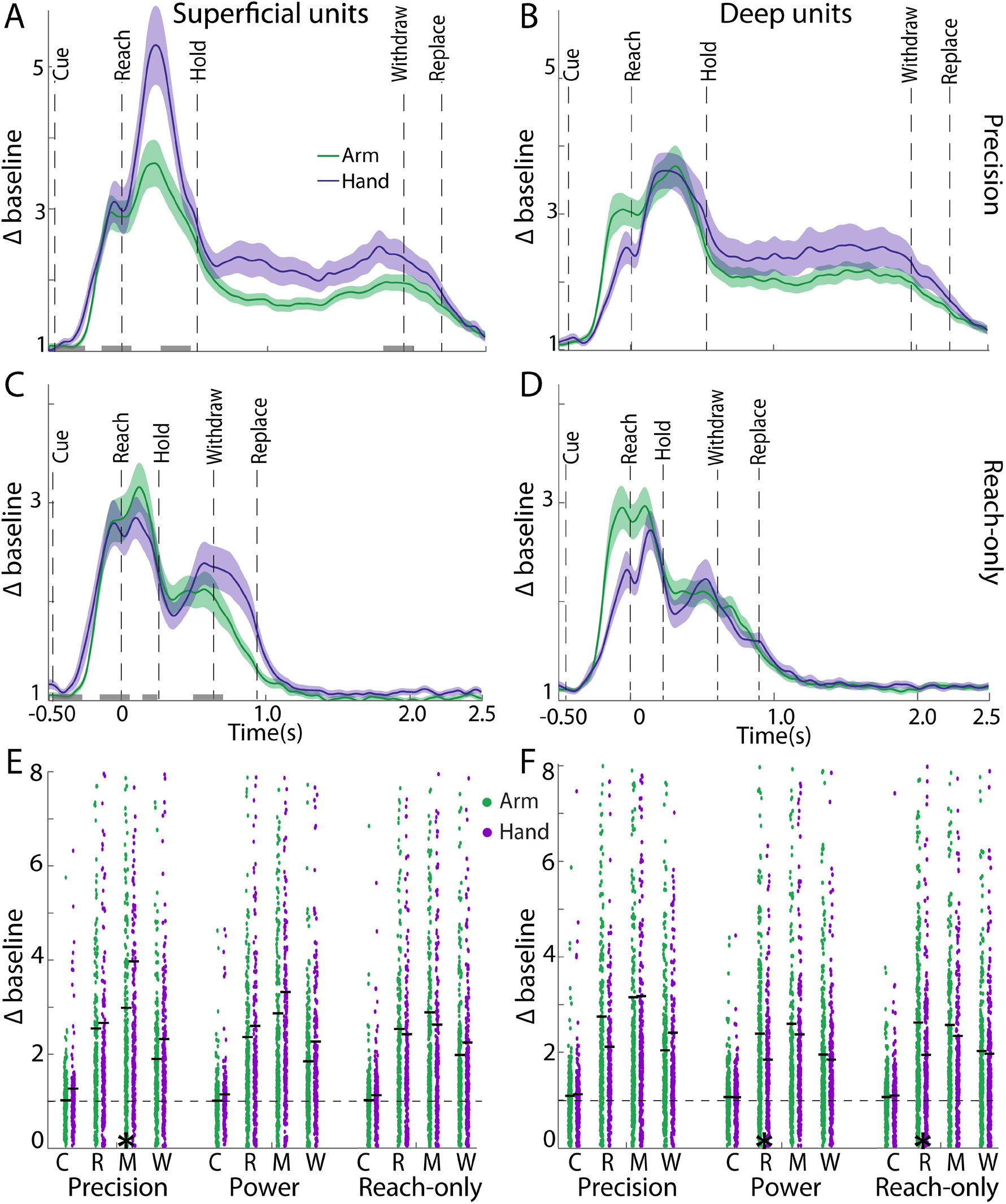
Higher neural activity in superficial laminar compartments. **(A)** Average PETHs for single units (n=295) recorded on the most superficial 16 channels (1 to 1600 mm from surface) of array penetrations in the M1 arm and hand zones. **(B)** Same as ***A***, but for single units (n=359) recorded on the deepest 16 channels (1700 to 3200 mm from surface). ***A-B*** are based on the precision condition. **(C-D)** Same as ***A-B***, but for the reach-only condition. **(E)** Distributions of superficial unit activity across conditions and task phases (C, R, M, W = cue, reach, manipulate, withdraw). Black horizontal line marks the mean of each distribution. Broken line is at baseline. (F) Same as ***E***, but for deep units. **\*** Post hoc difference (p<0.01) between arm and hand.

**Figure S3.**
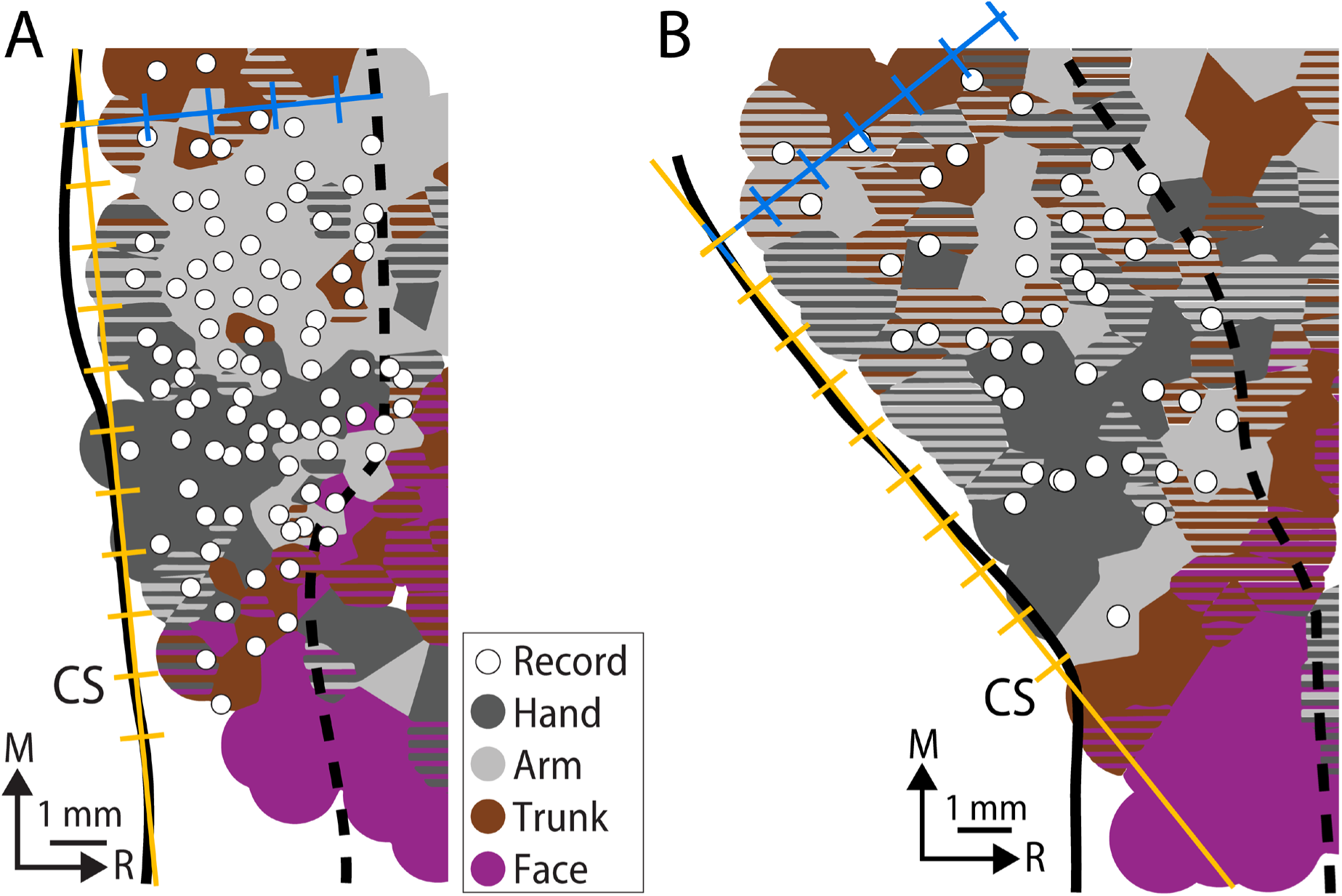
Standardizing recording coordinates across monkeys. **(A)** Recording sites superimposed on motor map (monkey G). The orange line is approximately aligned with the central sulcus (CS). Tick marks (1 mm) on the orange line reference distance in the medio-lateral axis with a zero point at the approximate border between the trunk and arm zones. Tick marks (1 mm) on the blue line reference distance in the rostro-caudal axis with a zero point at the central sulcus. **(B)** Same as ***A***, but for monkey S.

**Figure S4.**
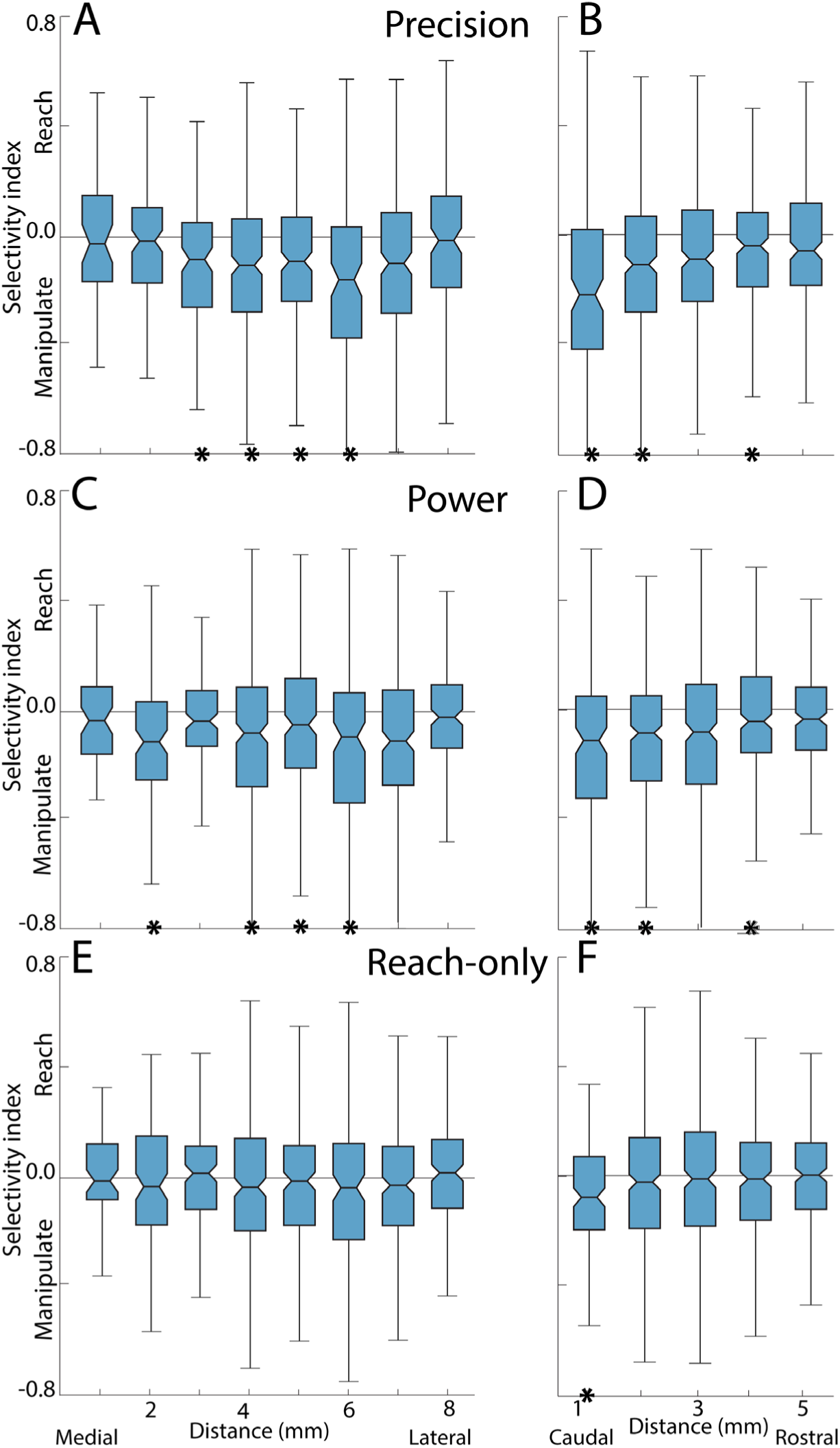
The spatial organization of task phase selectivity shifts with manual dexterity. **(A)** Selectivity index after binning units into 1 mm intervals in the medio-lateral axis. Data is from the precision grip condition in both monkeys. **\*** p<0.006 in one-sample t-test. **(B)** Same as ***A***, but binning here is in the rostro-caudal axis. **\*** p<0.01 in one-sample t-test. **(C-F)** Same as ***A-B***, but for the power and reach-only conditions. In all plots, dividing line is the median and box edges mark the 25^th^ and 75^th^ percentiles of the distribution. Whiskers span 1.5x interquartile range. In the precision and power conditions, there is significant selectivity for the manipulate phase in the center-caudal intervals, which overlap with the M1 hand zone. In contrast, selectivity is almost absent in the reach-only condition.

